# Learning sequence-function relationships with scalable, interpretable Gaussian processes

**DOI:** 10.1101/2025.08.15.670613

**Authors:** Juannan Zhou, Carlos Martí-Gómez, Samantha Petti, David M. McCandlish

## Abstract

Understanding the relationship between biological sequences, such as DNA, RNA or protein sequences, and their resulting phenotypes is one of the central goals of genetics. This task is complicated by epistasis, i.e., the context dependence of mutational effects. Advances in high-throughput phenotyping now make it possible to study these relationships at unprecedented scale, generating large datasets that measure phenotypes for tens or hundreds of thousands of sequences. However, standard regression models for analyzing such datasets often make unrealistic assumptions about the generalizability of mutational effects and epistatic coefficients across genetic backgrounds. Deep neural networks offer greater flexibility but suffer from limited interpretability and lack uncertainty quantification. Here, we introduce a family of interpretable Gaussian process models for sequence-function relationships that capture epistasis through flexible prior distributions that generalize classical theoretical models from the fitness landscape literature. In particular, these priors are parameterized by interpretable site-, allele-, and mutation-specific factors controlling the degree to which specific mutations decrease the predictability of the effects of other mutations. Using GPU acceleration to scale to large protein, RNA, and genome-wide SNP datasets, our models consistently deliver superior predictive performance while yielding interpretable parameters that both recover known features and uncover novel epistatic interactions. Overall, our methods provide new insights into the structure of the genotype-phenotype map and offer scalable, interpretable approaches for exploring complex genetic interactions across diverse biological systems.

## Introduction

Understanding how genetic sequences (DNA, RNA, and proteins) encode biological function remains one of the major open challenges in biology. Deciphering sequence-function relationships is crucial for understanding and predicting evolution [1–8] as well as for uncovering the genetic factors underlying human genetic and infectious diseases and cancer [9–11]. It is also essential for designing and engineering sequences with desired properties, such as maximizing the expression or activity of a protein of interest [12–14] or breeding crops and livestock with improved yield or tolerance to stress and pathogens [15–18]. Recent advances in high-throughput phenotyping technologies, such as deep mutational scanning and massively parallel reporter assays, have led to the rapid proliferation of datasets measuring functionality across large libraries of sequence variants [19]. These approaches have been applied to proteins [9, 20–37], regulatory sequences [38–48] and even genome-wide genetic variation [49–52], providing valuable insights into protein function and gene regulation. However, building models that capture the structure of these empirical sequence-function relationships and accurately predict phenotypes for novel genotypes remains challenging, in part because mutational effects often change due to the presence of other mutations [3, 5, 7, 8, 53–56].

Existing methods for modeling sequence-function relationships often make strong assumptions about the structure of this background dependence of mutational effects. For example, an additive model assumes that the effects of mutations are constant, such that the observed mutational effects in the data can be generalized to any novel background [57]. Pairwise interaction models relax this assumption by allowing mutational effects to vary, but still assume that the epistatic interaction between any pair of mutations is itself background-independent [58]. Perhaps surprisingly, this background independence of double-mutant epistatic coefficients implies that for any pairwise interaction model, the mean predicted phenotype changes quadratically as a function of the number of mutations from the wild type (or any arbitrarily chosen focal sequence)—a strong restriction on the types of geometric features such models can represent [55]. As a result, simple regression models often lack the flexibility to account for the full complexity of epistasis in empirical sequence-function relationships [55] and are frequently outperformed in prediction tasks by more expressive methods, such as neural networks [34, 35, 46, 59–66]. However, while neural networks are able to make more accurate predictions than traditional regression models, they provide limited capabilities for interpretability and uncertainty quantification, which reduces their utility.

Gaussian process models offer an alternative class of highly flexible models [55, 58, 67–74] that provide better interpretability and uncertainty quantification. These models work by defining a prior probability distribution over the space of possible sequence-function relationships and then computing the posterior distribution given the data [75], where in particular the prior distribution can be defined by interpretable parameters that control the predictability of mutational effects across different genetic backgrounds [55] and the posterior distribution provides a natural means to quantify uncertainty. While our previously described prior distributions for sequence-function relationships are isotropic, this is, they assume that every mutation has the same effect on the predictability of other mutations [55, 58, 70], empirical evidence suggests substantial heterogeneity. Different mutations can vary widely in how they affect predictability at other positions [28, 76] and in the total fraction of genetic variance explained by their interactions [77], with key mutations dramatically altering the effects of mutations elsewhere in the sequence [30, 77–79].

In this paper, we introduce a new family of anisotropic Gaussian process priors that capture heterogeneity in how mutations influence the predictability of other mutations based on two key principles. First, each position, allele, or mutation is assigned a decay factor that controls how much it affects the predictability of mutations at other positions. Second, when several mutations occur together, their effects on the predictability of other mutations are assumed to combine multiplicatively. These design choices enable us to generalize previously proposed theoretical models of fitness landscapes in several key directions [55, 77, 80], particularly by allowing us to generate families of complex random fitness landscapes using a moderately sized set of interpretable parameters, each tied directly to the behavior of individual mutations. We combine these new priors with recent advances in Gaussian process inference and modern GPU infrastructure [81–83] to analyze diverse high-throughput phenotyping experiments, including nearly complete genotype-phenotype maps for protein GB1 [84] and human 5*^′^*splice sites [42], a dataset for the longer AAV2 capsid protein [35], and a genome-wide dataset for yeast [50]. We then use our Gaussian process framework for a range of data-analytic tasks, including: (i) identifying sites, alleles, and mutations with strong effects on the predictability of other mutations, (ii) making phenotypic predictions for specific genotypes of interest and quantifying the uncertainty in these predictions, (iii) predicting the effects of all single point mutations across different genetic backgrounds and assessing our confidence in whether those effects have changed, and (iv) reconstructing combinatorially complete landscapes for moderately sized subsets of mutations. Overall, by studying these diverse systems, we find that mutations exhibit highly heterogeneous effects on the predictability of other mutations. By modeling this heterogeneity explicitly, we capture the background dependence of mutations with greater granularity and precision, thereby substantially enhancing the predictive power of our models.

## Results

### Gaussian processes for learning sequence-function relationships

To model sequence-function relationships, we use Gaussian process regression, which places a Gaussian prior distribution over the space of functions mapping genotypes to their phenotypes. This prior is fully specified by its mean and a covariance function, or kernel, *k*(*x, x^′^*), which defines the covariance of phenotypes between each possible pair of sequences *x* and *x^′^* under the prior. Gaussian process regression makes predictions based on the data *y* by computing the posterior mean for any genotype *x*

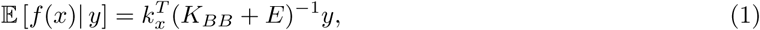

where *K_BB_*and *E* are square matrices modeling the prior covariance and noise variance for the set of measured genotypes *B*, and *k_x_* is the vector encoding the covariance between the genotype *x* whose phenotype we wish to predict and the set *B* of measured genotypes. A key advantage of Gaussian process regression is that it also provides a full characterization of the prediction uncertainty via the posterior covariance matrix (see Supplement 1.1).

Because Eq. 1 is a generic result applicable to all Gaussian process regression problems [75], the key determinant of model behavior and performance is the choice of the prior defined by the kernel function. In previous work, we introduced priors parameterized by biologically interpretable quantities such as the average squared epistatic coefficient [58, 70] or the variance associated with genetic interactions of each possible order [55]. These priors also control how the predictability of mutational effects decays with genetic distance [55].

In this section, we review our previously developed method, empirical variance component regression (VC regression) [55], propose three new methods that model how specific mutations influence the predictability of mutational effects at other sites, and analyze the flexibility in the set of expected variance components that can be under each prior.

### Empirical variance component regression

In [55], we introduced a family of flexible priors in which the kernel function for a pair of genotypes *x* and *x^′^* separated by *d* mutations is given by

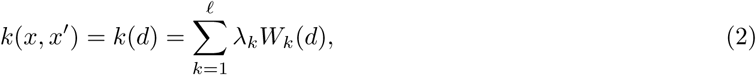

where *W_k_*(*d*) is a kernel that captures the covariance structure of pure *k*-th order interactions (given by a family of orthogonal polynomials known as the Krawtchouk polynomials, see [55, 85] for details) and *λ_k_ ≥* 0 is the expected variance of *k*-th order interactions under the prior. Importantly, this family of priors coincides with a well-studied class of theoretical random fitness landscapes where the covariance in phenotypes and mutational effects is completely determined by the Hamming distance between genotypes or genetic backgrounds [85–88]. These isotropic random landscapes are parameterized by the expected variance components of additive effects, pairwise, and each possible type of higher-order interaction between mutations. These variance components then control how fast the correlation in mutational effects between genetic backgrounds decays with the genetic distance between them.

Traditionally, the scalability of Gaussian process regression has been limited to fitting tens of thousands of training data points because computation of Eq. 1 and hyperparameter optimization scale cubically with dataset size [75, 81]. In the original proposal of VC regression [55], we inferred the model hyperparameters from the data by minimizing the discrepancy between the prior covariance and the empirical phenotypic covariance using weighted least squares [89] and exploited a sparse representation of the kernel matrix for the full sequence space to compute the posterior mean and covariances via iterative methods [55, 74]. This strategy allowed us to scale our methods to sequence spaces with up to 10 million sequences, equivalent to 12 nucleotides for DNAs/RNAs, 24 biallelic loci, or 5 sites for proteins.

Here, we alleviate these constraints on sequence length and alphabet size by leveraging recent developments in Gaussian process inference techniques and modern GPU computation using GPyTorch [81] and KeOps [83] to instead work with the dense matrix *K_BB_* (Eq. 1) whose dimensions depend on the size of the training data but not on the size of sequence space. These modern machine learning libraries implement algorithms that combine the power of GPUs and iterative methods to compute posterior means and covariances more efficiently, allowing computations for datasets with up to roughly 1 million observations [82]. Moreover, these techniques also enable a more principled approach for inferring hyperparameters, such as the *λ_k_*, via evidence maximization [75]. In other words, we find the combination of kernel hyperparameter values that maximize the likelihood of observing the empirical data (see Supplement 1.1). These improvements make our methods scalable to large datasets without constraints on the size of sequence space.

### Connectedness model

By tuning the relative contributions of different orders of epistatic interaction, VC regression allows priors that can reproduce any distance-dependent correlation function for mutational effects. However, the isotropic assumption that all mutations have equal effects on the predictability of other mutations still limits the expressivity of these priors. In this section, we introduce our first new model, which relaxes this isotropic assumption by allowing mutations at each site to exert a site-specific characteristic effect on the predictability of the phenotypic effects of mutations at other sites.

To construct the model, we introduce a site-specific decay factor *δ_p_* for each of the *ℓ* positions (*p* = 1*, . . ., ℓ*). Biologically, *δ_p_* captures how much a mutation at site *p* disrupts the predictability of mutational effects across genetic backgrounds. For example, if two genotypes differ only at position *p*, then the correlation under the prior in the effects of mutations at all other positions is precisely 1 *− δ_p_*. A small *δ_p_* suggests that mutations at *p* have minimal effects on the effects other mutations, while a large *δ_p_* indicates strong epistatic influence. To generalize this idea to genetic backgrounds differing at multiple sites, we assume that the effects of different mutations on the predictability of other mutations combine multiplicatively. That is, mutating an additional position *q* further reduces the original correlation by a factor of 1 *− δ_q_*, producing an expected correlation of (1 *− δ_p_*)(1 *− δ_q_*). This results in a biologically interpretable kernel function, where the covariance between two sequences decreases according to the cumulative effect of their mutational differences:

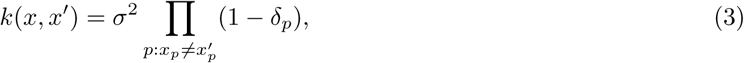

where *σ*^2^ represents the shared variance across all genotypes. Importantly, the decay factors *δ_p_* not only control the phenotypic correlation between different sequences, but also the correlation in mutational effects in genetic backgrounds that differ at specific positions *p*, given by *_p_*(1 *− δ_p_*) (see Supplement 1.2).

Our prior is closely related to the ‘connectedness model’ [77], a statistical model of Gaussian fitness landscapes first proposed for studying evolution of the distribution of fitness effects for adapting populations. The connectedness model is parameterized by *ℓ* site-specific probabilities 0 *≤ µ_p_ ≤* 1, corresponding to the tendency for each site *p* to be involved in epistatic interactions. In particular, the strength of interactions among any set of positions is proportional to the product of their corresponding *µ_p_* factors.

In Supplement 1.4, we show that Eq. 3 provides a slight generalization of the connectedness model and that the two site-specific factors *µ_p_*and *δ_p_*are related through the following equation:

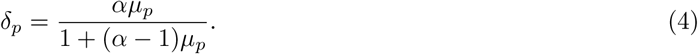

Interestingly, we show that our connectedness kernel in Eq. 3 remains a valid (i.e. positive-definite) kernel for some *δ_p_ >* 1, which would correspond to probabilities *µ_p_ >* 1 in the original connectedness model. In such cases, introducing a mutation with a decay factor *δ_p_* larger than one leads to anticorrelated phenotypes and mutational effects. This added flexibility enables the model to capture a broader range of epistatic patterns than the original connectedness model, although empirical support for this so-called egg box [28, 76] correlation structure remains limited. In addition, while the original connectedness model was proposed for biallelic genotypic spaces, our connectedness kernel is compatible with an arbitrary number of possible alleles per site.

The relationship between the connectedness kernel and the VC regression kernel can be clarified by examining how each generalizes a simple kernel where the correlation between genotypes decays geometrically with Hamming distance *d*(*x, x^′^*), i.e. *k*(*x, x^′^*) = *σ*^2^*β^−d^*^(^*^x,x′^*^)^. This geometric decay kernel includes the exponential kernel as a special case (*β >* 0). The exponential kernel itself arises when applying the standard radial basis function (RBF) kernel to one-hot encoded sequences [67, 87], when the magnitude of epistatic interactions decays exponentially with interaction order (i.e. *λ_k_* = *e^−b×k^*) [55, 73, 87] and as the diffusion kernel on the Hamming graph [69, 90]. Our VC regression kernel can then be viewed as a generalization of the geometric kernel to allow the *λ_k_* to deviate from this strict multiplicative scaling while retaining the isotropic property of the geometric kernel. In contrast, the connectedness kernel offers a different generalization: whereas the geometric decay kernel treats all mutations the same (*β* = (1 *− δ*) and *δ_p_* = *δ*), the connectedness kernel introduces a detailed dependence on the mutated sites through site-specific decay factors (or equivalently, site-specific epistatic strengths). Interestingly, both the VC regression and the connectedness priors have the same number of hyperparameters so that comparing their performance provides an indication as to the relative importance of precisely matching the contribution of higher-order epistasis versus incorporating site-to-site heterogeneity in the extent of epistasis.

### Jenga model

The connectedness model introduces flexibility to the prior by allowing the extent to which mutational effects generalize across genetic backgrounds to depend on which specific sites are mutated. One limitation of this approach is that it treats all mutations at a given site equally. In practice, however, some alleles may be more likely than others to participate in genetic interactions and to influence the predictability of mutations at other sites. In this section, we generalize the connectedness model by allowing the decay in the correlation of mutational effects to depend not only on the mutated sites, but also on which specific alleles have been altered.

In the connectedness model, a mutation at site *p* reduces the correlation in mutational effects at other sites by a factor of 1 *− δ_p_*. Here, instead of assigning a uniform decay factor *δ_p_* to all mutations at site *p*, we assign an allele-specific decay factor *δ^a^*, such that mutating allele *a* to a new allele *a^′^* results in an overall reduction in correlation by 100 *×* (1 *− δ^a^*)(1 *− ′*percent. Assuming that the effects of mutations on phenotypic correlations across positions combine multiplicatively in this manner, we obtain a new distribution over the space of all sequence-function relationships, with covariance characterized by the following kernel function:

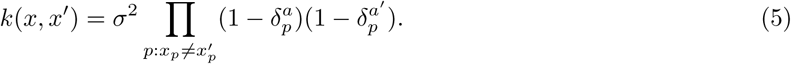

Importantly, this expression also generalizes to describe the decay in the predictability of mutational effects, such that the correlation in these local effects is IT (1 − δa)(1 − a′, for all positions *p* segregating between It is easy to see that this model contains the connectedness model as a special case when all alleles at a given position have a common decay factor. Furthermore, in the special case *δ^a^* = 1, according to Eq. 5, mutating allele *a* results in zero correlation in phenotypes or mutational effects. Indeed, if all alleles in Eq. 5 have decay factors equal to 1, we recover the classical ‘House-of-Cards’ (HoC) model [80], in which mutating any allele completely eliminates predictability in phenotypes or mutational effects—akin to how removing any component from a house made of playing cards causes the entire structure to collapse. Thus, by allowing alleles to have a continuum of effects on the overall correlational structure, our model can be viewed as a generalization of the HoC model where different alleles at a site have different effects on the correlation structure. We therefore refer to this new model of random fitness landscapes as the ‘Jenga model’, reflecting the idea that some mutations may have negligible effects, while others can disrupt the entire correlational structure, much like how specific blocks in the game Jenga can either be easily removed or cause the whole tower to fall.

The Jenga kernel is also closely related to the classical automatic relevance determination (ARD) kernel [75, 91] applied to the one-hot encoding of genotypes (see Supplement 1.5). However, one important difference is that the allele-specific decay factors *δ^a^*in the Jenga model are allowed to exceed one under conditions that ensure the positive definiteness of the kernel on sequence space as a whole (see Supplement 1.6). As a result, whereas the ARD kernel requires all correlations to be strictly positive, the Jenga kernel permits negative correlations between phenotypes and mutational effects across adjacent genetic backgrounds, thereby offering the flexibility to model eggbox-like patterns that could, in principle, be induced by mutations at some positions.

### General product model

In the Jenga model, the effect of a mutation on the correlation between genotypes is parameterized by multiplying the two allele-specific factors, meaning that each allele makes the same contribution to the decay in correlations for all mutations involving that allele. In this section, we relax this assumption to further generalize the Jenga model. We do so by assigning a mutation-specific decay factor *δ^a,a^* to each pair of alleles *a, a^′^*, such that mutating from *a* to *a^′^* (or vice versa) reduces the overall correlation by a factor of (1 *− δ^a,a′^*).

This yields the following general product kernel:

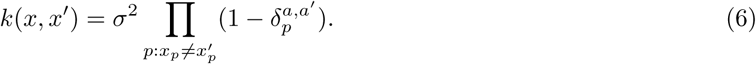

Similar to the Jenga model and connectedness model, the correlation in mutational effects is given by the product of the mutation-specific decay factors for sites segregating between the two genetic backgrounds: IT (1 − δa,a′).

It is easy to see that Eq. 6 provides greater flexibility than the Jenga model, as the general product kernel has *^α^* hyperparameters per site, whereas the Jenga model has only *α* hyperparameters. In fact, Eq. 6 represents the most general form of a homoscedastic product kernel such that any product kernel with uniform variance can be expressed in this form and can be guaranteed to remain positive definite through an appropriate parameterization [92].

### Variance components of the connectedness, Jenga, and general product kernels

We can understand certain aspects of the geometry of random fitness landscapes such as the connectedness model, the Jenga model, and the general product model by quantifying how much of the overall phenotypic variance is expected to be explained by epistatic interactions of different orders under these priors. While VC priors are directly parameterized in terms of these expected variance components, our new random fitness landscape models are parameterized by decay factors, which have a less straightforward relationship to variance components.

In the Supplement 1.7, using a general result for deriving variance components from arbitrary product kernels, we show that all three models are capable of capturing epistatic interactions of every possible order between any number of positions. Furthermore, we find that a single decay factor affects all expected variance components, such that increasing any decay factor systematically shifts the variance toward higher-order components.

We also find that there are certain combinations of expected variance components that cannot be obtained by tuning the decay factors of our three new priors, for example, priors with interactions of a single order alone (e.g., additive, pairwise). This behavior is more limited than in the VC kernel, which can match any valid combination of expected variance components. Furthermore, we show that the set of expected variance components under a prior from the connectedness model is in fact the same as that of based on the general product model (see Supplement 1.7, Theorem 3). Therefore, although the connectedness model has the fewest hyperparameters, it is just as expressive as the general product and Jenga models in terms of realizable combinations of expected variance components when used as prior distributions. Finally, although these new priors can express only limited set of variance components in expectation, it is important to realize that the posteriors can still capture arbitrary patterns of epistasis given sufficient data, since these priors produce a positive density for any possible landscape.

### Applications to experimental genotype-phenotype datasets

In the previous section, we introduced new models of random sequence-function relationships in which the decay in the predictability of mutational effects between genetic backgrounds is determined by the positions, alleles, or mutations at which sequences differ. Importantly, the connectedness model, the Jenga model, and the general product model form a hierarchical family, with each representing a generalization of the preceding model. These kernels equip us with the flexibility to model complex epistasis in the data while allowing for a balance between model expressivity, interpretability, and computational efficiency.

In this section, we implement these models as prior distributions in a Gaussian process framework for modeling empirical genotype-phenotype datasets. We infer the site-, allele-, or mutation-specific decay factors from the data by fitting the kernels in Eq. 3, Eq. 5, and Eq. 6 through evidence maximization, and we use them for two main purposes. First, the inferred decay factors provide important broad-scale insights into the structure of epistasis in the empirical sequence-function relationship by highlighting mutations that reduce the predictability of other mutations. Second, we use the inferred decay rates as hyperparameters of a prior distribution for Gaussian process regression to predict phenotypes for unobserved genotypes and to assess our confidence in these predictions.

### Application to protein GB1

We first applied our methods to the protein GB1 dataset [84], which contains relative enrichment values of nearly all 20^4^ = 160, 000 genotypes across four highly epistatic amino acid sites on the IgG-binding domain of streptococcal protein G, where selection was imposed for IgG-binding.

To assess the predictive performance of our methods, we first fit several standard models for studying sequence-function relationships to this dataset, including a simple additive model, a pairwise interaction model, and a global epistasis model, which maps the sequence first to an underlying additive phenotype and then onto the observation scale through a nonlinear monotonic function [61, 94, 95]. We also fit Gaussian process regression models with the VC kernel and the geometric kernel, with the model hyperparameters optimized via evidence maximization. The out-of-sample performance of these models under different proportions of training data is summarized in Figure 1A. We first found that the additive model achieves an *R*^2^ of approximately 0.5 on the test data, and that adding a global nonlinear function to the additive phenotype increased the test *R*^2^ to slightly over 0.65. Adding pairwise interaction terms to the additive model without a global epistasis nonlinearity resulted in a more substantial improvement, with test *R*^2^ values close to 0.75 when the training data proportion exceeded 10%. Next, we found that Gaussian process regression using the VC kernel and geometric kernel show similar performance, achieving a test *R*^2^ over 0.8 at high data density, corresponding to a 5% improvement over the pairwise model.

**Figure 1:**
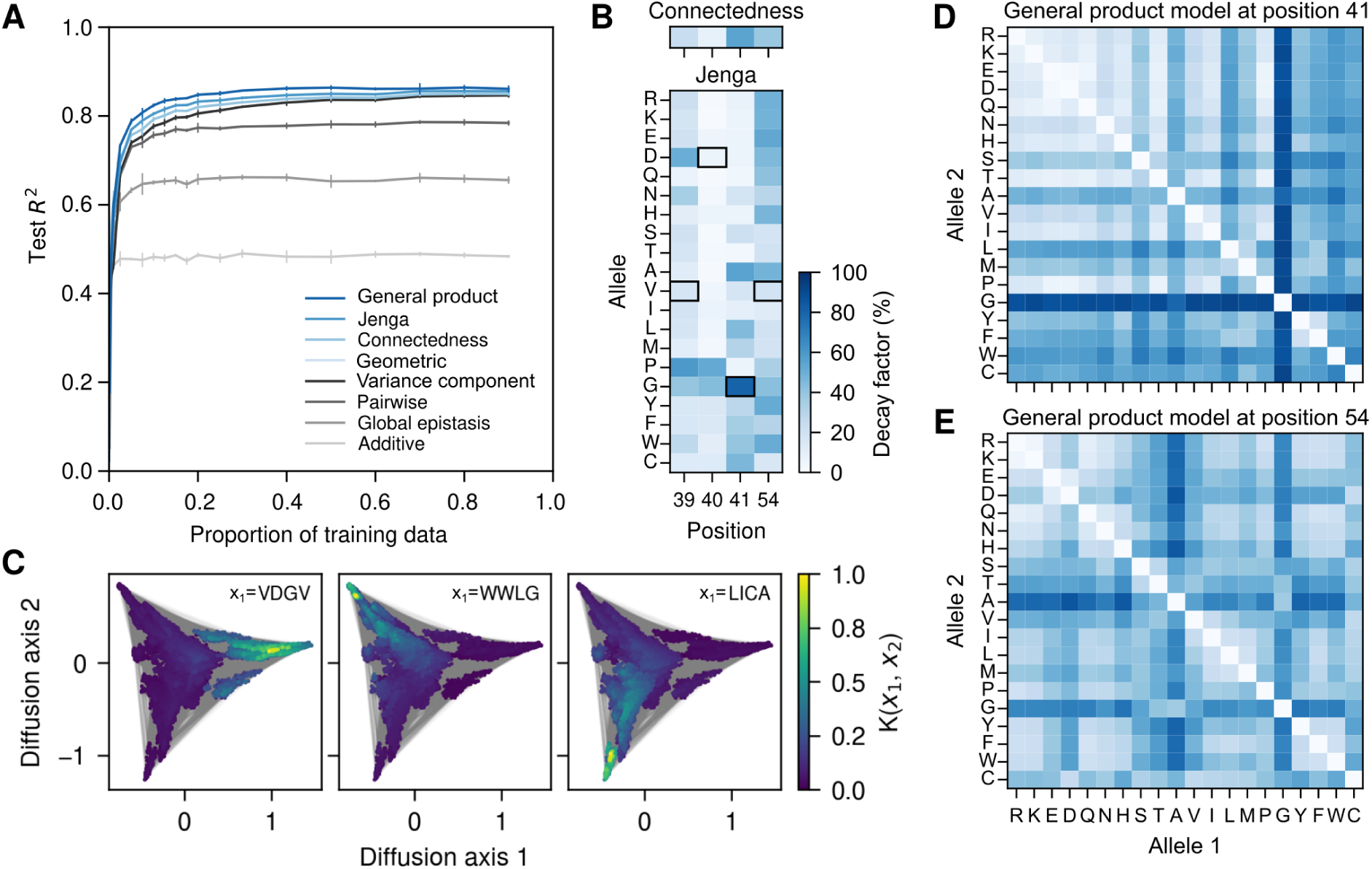
Application to the genotype-phenotype dataset for protein GB1. (A) Model predictive performance, measured by the *R*^2^ on held-out data, as a function of the fraction of data used for training. Error bars represent one standard deviation across three independent subsets of sequences used for training at each proportion. (B) Inferred position- and allele-specific decay factors under the connectedness model (top) and Jenga model (bottom), respectively. Black squares highlight the allele of the wild-type sequence at each position. (C) The Jenga prior is able to capture information about the three main fitness peaks of the GB1 landscape [58]. In each panel, dots represent genotypes and lines denote single point mutations between genotypes. Squared distances between dots optimally approximate the commute times between genotypes under a weak-mutation evolutionary model under selection for high phenotypic values [74, 93] (see Methods). Genotypes are colored by their correlations with three genotypes VDGV, WWLG and LICA, each representing a local fitness peak. Correlations were calculated using the Jenga kernel with hyperparameters inferred using evidence maximization. The complete GB1 fitness landscape for the evolutionary model was constructed using the maximum a posteriori estimate under the Jenga model. (D, E) Inferred mutation-specific decay factors under the general product model for positions 41 and 54. These matrices show great heterogeneity in the influence of mutations at a given site on the predictability of mutational effects at other sites both in terms of the degree of influence conferred by mutations between different pairs of alleles and in the effect of mutations between the same pair of alleles at different sites.

With the performance of these baseline methods established, we proceeded to fit our three new methods to quantify site-, allele-, or mutation-specific effects on the predictability of other mutations and examine how accounting for heterogeneity at various levels of the correlational structure improves predictive performance. First, we found that the connectedness model indeed reveals heterogeneous effects across the four amino acid positions. Specifically, we found the site-specific decay factors inferred using evidence maximization showed a nearly seven-fold difference (*δ*_39_ = 0.23, *δ*_40_ = 0.08, *δ*_41_ = 0.54, *δ*_54_ = 0.38; Figure 1B). This suggests that mutating sites 41 and 54 have the strongest effect on the predictability of phenotypes and mutational effects, such that the same mutations have an average correlation of 0.53 across backgrounds segregating only at site 41 or 54, and an average correlation of 0.28 when both sites segregate (see Eq. 3). In contrast, mutations at site 40 have a much smaller effect on the predictability of other mutations, such that for genetic backgrounds segregating only at site 40 the same mutations have an average correlation of 0.92. This rank order between sites is consistent with previous analysis showing that higher-order interactions in this dataset are enriched among site 39, 41 and 54, due to steric interactions [23, 84]. We found that using these decay factors to parameterize the connectedness kernel for Gaussian process regression yields a slight improvement in predictive accuracy over both VC regression and geometric kernel regression (Figure 1A).

We next fit the Jenga model to the data and examined how specific alleles at each position affect the correlation in phenotype and mutational effects. We first present the inferred allele-specific decay factors as a heatmap in Figure 1B. The decay factors exhibit substantial heterogeneity across alleles. For example, at position 40, mutations involving Gly and Pro result in more than a 50% reduction in correlation, whereas mutations involving other amino acids have minimal or negligible effects. The Jenga model heatmap is also consistent with the aforementioned steric interaction model [23, 84]. This is most evident at position 41, where amino acids with the highest decay factors tend to have either small side chains (e.g., Gly and Ala) or bulky ones (e.g., Leu, Trp, and Phe). These mutations likely cause the largest disruption to the optimal pattern of steric interactions, possibly resulting in nonfunctional proteins and consequently a breakdown in the correlation of mutational effects. Finally, we observed a modest improvement in *R*^2^ for the Jenga model over the connectedness model, which was comparable in magnitude to the improvement of the connectedness model over VC regression (Figure 1A).

To gain some intuition as to how Jenga regression works, we examined the qualitative structure of the Jenga kernel using a visualization technique to understand how the correlations incorporated into the Jenga kernel relate to the structure of the genotype-phenotype map. In Figure 1C, we first represented all 160,000 genotypes in the GB1 fitness landscape using a 2-dimensional embedding generated using a method similar to diffusion maps[74, 93]. The method is based on modeling how a population would evolve across this fitness landscape, and results in a triangular structure with three high-fitness clusters, i.e. fitness peaks, occupying the corners and separated by a central mass of low-fitness genotypes. Importantly, we previously found that the local structure within each cluster is mainly additive such that the effects of mutations are well-correlated across backgrounds within clusters but not between clusters [58, 96]. We next chose a focal sequence from each of the three clusters and colored each sequence by its correlation with this focal sequence under the Jenga prior (Figure 1C). We observe that all three high-fitness genotypes exhibit strong correlation with neighboring genotypes within the same cluster, and that this correlation decays to near zero outside each cluster. This indicates that the model predominantly uses mutational effects observed within the same cluster to make predictions for a focal genotype, and thereby in essence is fitting multiple independent models across different regions of the fitness landscape. For comparison, we also present the same visualization for all models fit to the GB1 data in Figure S1. Unlike the highly localized correlation structure of the Jenga prior, the isotropic priors (which only depend on the number of mutations separating two sequences and not their identity, e.g. the VC and geometric priors) show strong to moderate correlation between each focal genotype and genotypes across the whole landscape. The key reason for this qualitative difference is that the geometric kernel can only capture the typical or average effect of mutations on the predictability of other mutations. Thus, two genotypes that are mutationally adjacent will still have a high correlation even if the mutation that separates them is particularly epistatic. In contrast, the Jenga model’s ability to account for variation in epistatic effects across alleles enables it to model the lack of correlation between adjacent genotypes when separated by strong-effect mutations, while allowing distant genotypes to maintain high correlation if separated only by mutations with small decay factors. As a result, the Jenga kernel more closely reflects the structure of the GB1 landscape, which likely contributes to its improved predictive accuracy.

Finally, we fit our most flexible model, i.e., the general product model. Unlike the Jenga model, which assigns decay factors to individual alleles at each position, the general product model directly characterizes the effects of all mutations. This added flexibility yields another modest improvement in predictive accuracy, comparable in magnitude to the improvement achieved by the Jenga model over VC regression (Figure 1A). To investigate the cause of this improvement, we visualized the mutation-specific decay factors as 20 *×* 20 matrices. We first examined the matrices for positions 41 and 54 in Figures 1D and 1E, and then considered side-by-side comparisons of the matrices inferred by the general product model with those constructed by multiplying the corresponding allele-specific decay factors from the Jenga model (Figure S2). We found that the effects of some mutations in the general product prior can sometimes be well approximated by the Jenga prior; for example, mutating Gly at position 41 uniformly induces strong epistasis in both the Jenga model and the general product priors. However, there are also mutations with mutation specific decay factors that deviate from the Jenga prior’s expectation. For example, according to Figure 1E, the three aromatic residues Tyr, Phe, and Trp at position 54 are largely interchangeable, but under the Jenga prior, mutations among these residues are expected to have moderately large epistatic effects on other mutations (Figure S2). This difference arises because the Jenga prior can only encode the average epistatic effects of each allele, whereas the actual effects of mutations at position 54 have a more subtle dependence on the physical properties of the specific alleles involved.

### Application to human **5*^′^*** splice site

Having demonstrated the performance of our methods on the protein GB1 dataset, we now apply our methods to modeling an RNA sequence-function relationship. The experimental data consists of the activity of nearly all variants covering 8 nucleotides on the 5*^′^* splice site of exon 7 for the survival of motor neuron 1 (SMN1) gene in a minigene construct [42], with splicing activity measured as the proportion of correctly spliced transcripts (percent spliced in, PSI).

To generate performance baselines, we first fit the additive, pairwise epistatic, and global epistasis models to the data. In Figure 2A, we can see that the additive model exhibits poor predictive accuracy, with test *R*^2^ around 0.15. The pairwise model provides a substantial improvement, achieving an *R*^2^ of 0.45 when the training data proportion exceeds 20%. However, the pairwise model is outperformed by the global epistasis model, which typically achieved an *R*^2^ of approximately 0.6. This rank order is consistent with our previous understanding of the SMN1 splicing landscape, which at a coarse scale can be approximated as a single-peaked fitness landscape resulting from an additive fitness landscape transformed by a steep sigmoidal nonlinearity on the measurement scale (Figure S3A) [55].

**Figure 2:**
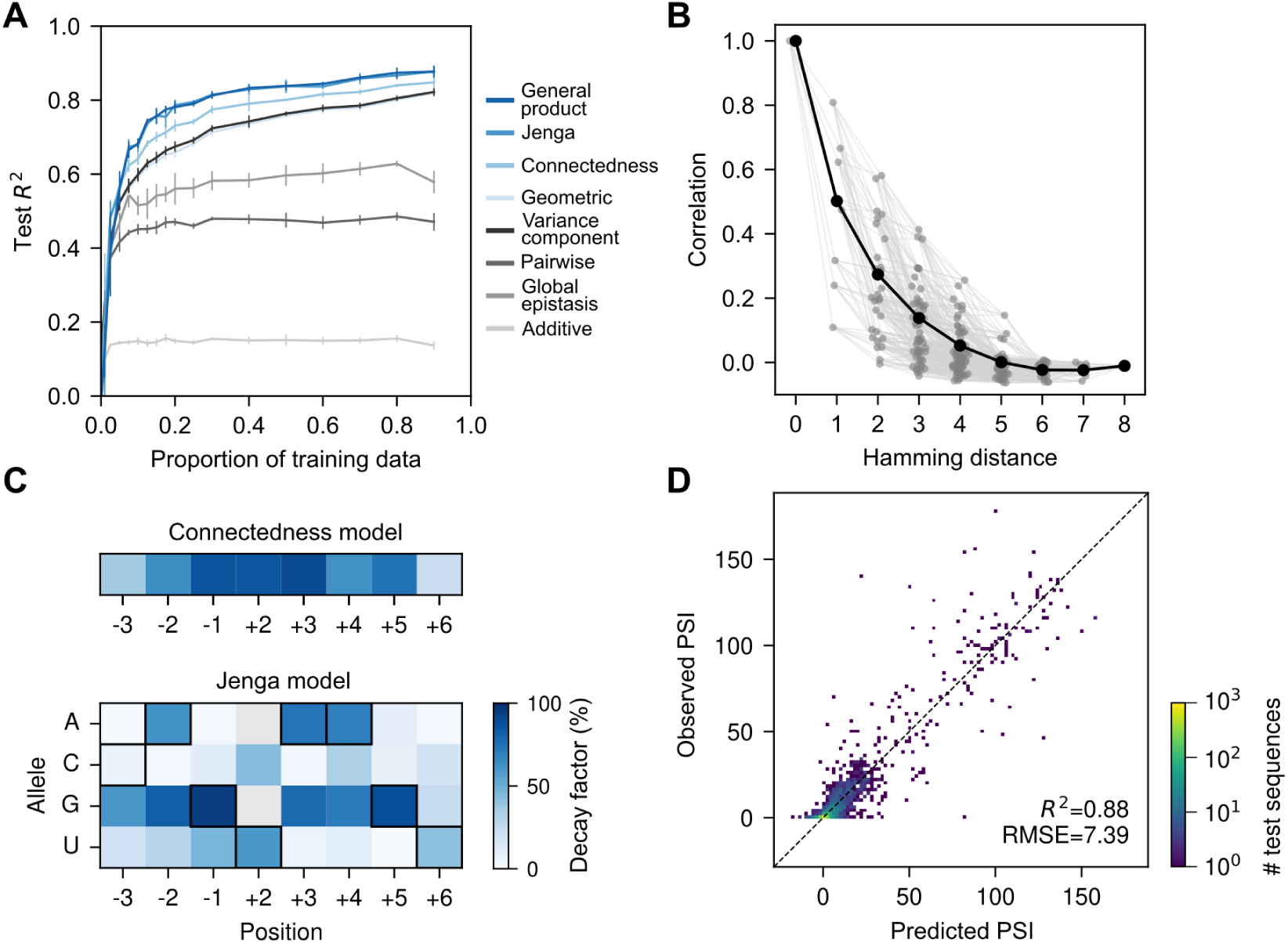
Application to the high-throughput dataset for 5*^′^*splice site of SMN1 exon 7. (A) Model predictive performance, measured by the *R*^2^ on held-out data, as a function of the fraction of data used for training. Error bars represent one standard deviation across three independent subsets of sequences used for training at each proportion. (B) Empirical correlations between pairs of sequences. Black dots show the average correlation for all pairs of sequences at a given Hamming distance. Gray dots show the correlation between genotypes segregating at specific sets of positions for each distance class. Values on the x-axis were jittered to facilitate visualization. (C) Inferred position- and allele-specific decay factors under the connectedness model (top) and Jenga model (bottom), respectively. Black squares highlight the canonical 5*^′^* splice site nucleotides complementary to the U1 snRNA template. (D) Two-dimensional histogram comparing the observed Percent Spliced In (PSI) values against predictions of the Jenga model on test 5*^′^* splice sites, comprising 10% of the measured genotypes.

The limited predictive performance of these basic models suggest that the data contain a more complex pattern of genetic interactions. In fact, the empirical distance-correlation function (Figure 2B, black dots) shows that correlation between pairs of sequences in the data cannot be accurately modeled by a linear or quadratic function as expected under an additive or pairwise interaction model [55]. Instead, each mutation, in average, seemed to decrease the correlation by about half, as expected under a geometric kernel. In line with this, we found that Gaussian process regression with the VC kernel and the geometric kernel showed similar performance throughout the range of training data density, with test *R*^2^ increasing nearly linearly once the training data proportion exceeded 20%, and surpassing 0.8 with dense training data. This improvement over the global epistasis model can be attributed to their ability to fit higher-order epistasis, which can provide a good approximation of the global nonlinearity while also capturing mutation-specific interaction patterns, often referred to as specific epistasis [97], that the global epistasis model cannot capture.

However, when stratifying the empirical distance-correlation function depending on the sites at which sequences differ (Figure 2B, gray dots), we can see that the empirical correlations exhibit large variation within each distance class, to the extent that two genotypes differing at up to five positions can be more correlated than pairs separated by only one position. This observation suggests that mutations at different sites in the SMN1 dataset have very heterogeneous effects on the predictability of other mutations. Consistent with this observation, the connectedness model yielded an improvement in predictive performance, typically increasing test *R*^2^ by 3–4% over the VC and geometric kernels (Figure 2A). In Figure 2C, we show the decay factors of the connectedness prior inferred using evidence maximization. We can see that some positions when mutated can reduce the correlation under the prior by over 80% (e.g. *δ_−_*_1_ = 0.86, *δ*_+2_ = 0.85, *δ*_+3_ = 0.89), while others have more moderate effects on the correlation (*δ_−_*_3_ = 0.36, *δ*_+6_ = 0.23). This allows the connectedness prior to better capture the empirical correlation structure observed in Figure 2B than the VC regression prior, which depends on mutational distance but not the specific sites at which sequences differ.

Next, we turn to the Jenga model to examine how changing specific alleles affects the predictability of other mutations. We find that the allele-specific decay factors of the Jenga model inferred using evidence maximization show substantial variation among alleles within each site (Figure 2C). Comparing the decay factors of the connectedness model and the Jenga model, we find that the former provides a coarse approximation of the latter, such that sites with large allele-specific decay factors also tend to have large site-specific decay factors under the inferred connectedness prior. On the other hand, the additional flexibility of the Jenga model is reflected in a roughly 3% improvement in *R*^2^ over the connectedness model at most training data proportions.

Interestingly, we find that the alleles with the largest decay factors under the Jenga model often coincide with the most preferred nucleotides under the global epistasis model (Figure S3B), which infers the presence of a sharp, threshold-like effect, consistent with the biophysical intuition that PSI can be expressed as a sigmoid function of the binding affinity between the 5*^′^* splice site and the U1 snRNA [43, 55, 98]. Because mutating critical nucleotides (e.g., U at position +2) typically results in nonfunctional splice sites, subsequent mutations have little to no effect since the splice site has already been rendered inactive. Consequently, these critical alleles are inferred to have large decay factors in the Jenga model, as mutational effects in backgrounds with and without the allele tend to be poorly correlated. This ability to incorporate different phenotypic correlations for different mutations appears to result in a qualitatively better fit of the global nonlinearity, so that while VC regression produces a systematic nonlinear pattern of residuals (Figure S3C), inference under the Jenga model produces a pattern of residuals for out-of-sample predictions centered on the observed values (Figure 2D). Overall, these results show that beyond capturing specific epistatic interactions, our new models continue to perform well in the presence of a strong global nonlinearity.

Finally, we applied our most flexible model, general product regression, to the data. Unlike in the protein GB1 example, allowing mutation-specific decay factors for SMN1 does not improve predictive performance relative to the Jenga model (Figure 2A). Examination of the inferred mutation-specific decay factors shows strong agreement with those implied by the Jenga model (Figure S3D), which accounts for their nearly identical performance. A contributing factor to the differing behavior of these models across datasets may be the alphabet size, as the difference in parameter count and expressivity between the Jenga and general product models is much smaller in nucleotide space (4 alleles versus 6 allele pairs) than in amino acid space (20 alleles versus 190 allele pairs).

### Application to AAV2 capsid protein

So far, we have applied our methods to relatively small sequence spaces. To demonstrate their utility in larger sequence spaces, we now apply them to a sequence–function dataset for the capsid protein of adeno-associated virus 2 (AAV2). This dataset encompasses 42,328 variants targeting 28 amino acid sites (positions 561–588), spanning buried, surface, and interface regions [35, 99], with corresponding deep mutational scanning (DMS) scores measuring viral production efficiency. Importantly, the sequence space for this example contains 20^28^ *≈* 2.7 *×* 10^36^ sequences, far larger than could be accommodated by our previous approaches [55, 58, 70, 74] which could only handle sequences spaces containing low millions of sequences.

As in previous sections, we first fit the baseline additive and global epistasis models across increasing proportions of training data (Figure 3A). Both models exhibit strong predictive performance, achieving test *R*^2^ values close to 0.6 and 0.8, respectively, as the training data proportion increases. The pairwise interaction model performs comparably to the global epistasis model at higher sampling densities but is inferior at lower sampling densities. We next fit the geometric, connectedness, Jenga, and general product models to the AAV2 dataset, selecting model hyperparameters via evidence maximization. We did not explicitly fit the VC regression to this dataset because the dataset lacks pairs of sequences at all relevant distances, preventing reliable estimation of the empirical autocorrelation function. The more expressive priors (Jenga and general product models) consistently outperformed the geometric and connectedness priors when trained with over 30% of the data. In contrast, the simpler priors performed better when less training data was available, a pattern that differs from our earlier results using nearly complete combinatorial data for short sequences. This difference is expected, as this dataset samples only a small fraction of the possible sequence space, and the number of hyperparameters to learn varies by an order of magnitude across models. For example, the general product prior has 5,322 hyperparameters, compared to 562 for the Jenga prior and only 28 for the connectedness prior.

**Figure 3:**
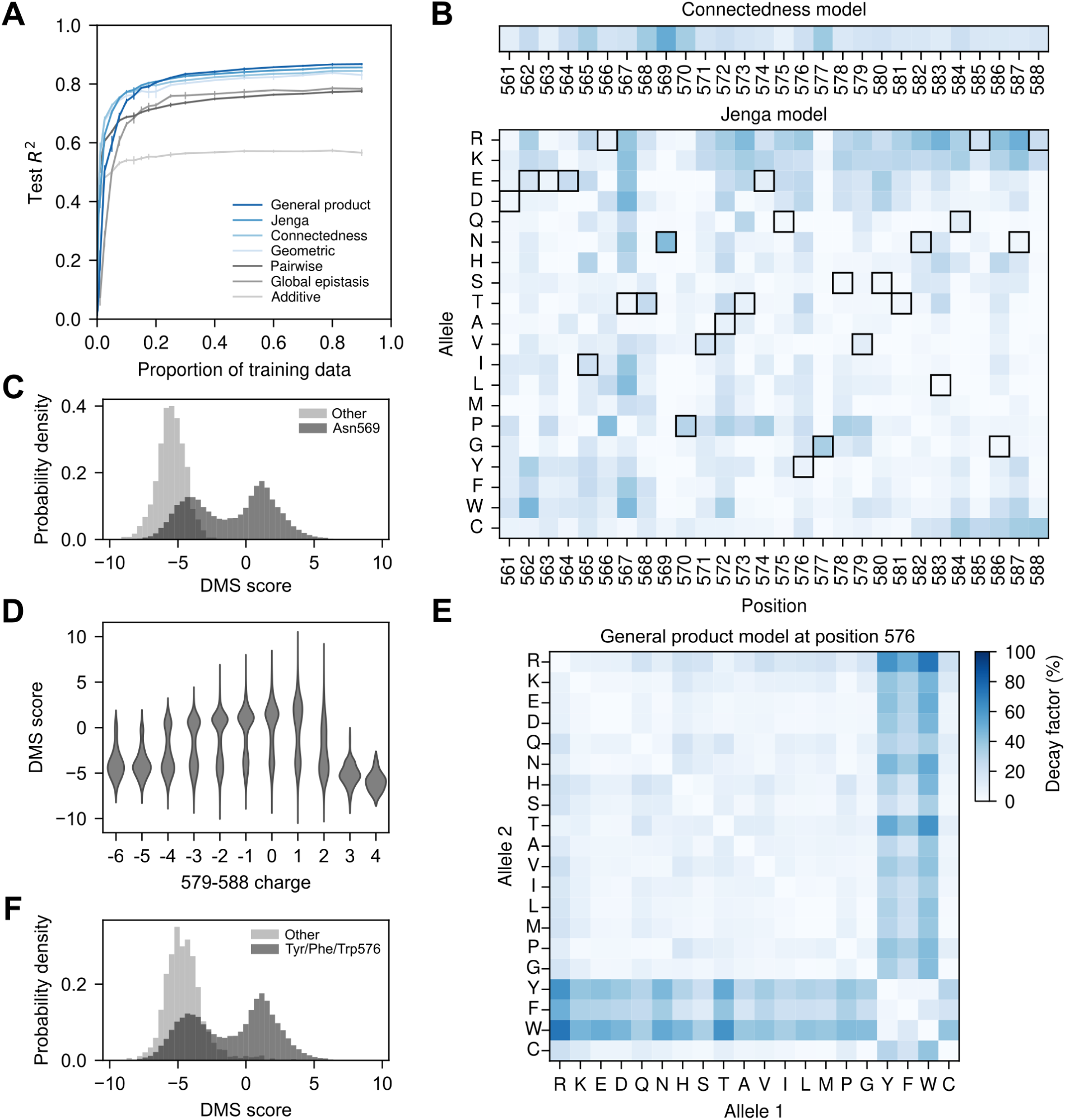
Application to the AAV2 capsid protein. (A) Model predictive performance measured by test *R*^2^ on held-out data, shown as a function of the proportion of training data. Error bars represent one standard deviation across three random subsets of training sequences at each proportion. (B) Inferred decay factors for each of the 28 positions under the connectedness model (top) and for 20 amino acids at each position under the Jenga model (bottom). Black squares indicate the wild-type allele at each position. (C) Distribution of raw DMS scores for sequences with Asn at position 569 (dark gray) compared to sequences with other amino acids at that site (light gray). (D) Raw DMS scores as a function of net charge across amino acids 579–588, indicating that intermediate charge levels are associated with higher viral production. (E) Inferred mutation-specific decay factors from the general product model for position 576. (F) Distribution of raw DMS scores for sequences with an aromatic residue (Tyr, Phe, or Trp) at position 576 (dark gray) compared to sequences with other amino acids at that site (light gray).

In the previous examples, we found that the site-, allele-, and mutation-specific decay factors are often related to the qualitative properties of complex sequence–function relationships and can help identify regions of sequence space where mutational effects differ systematically. We next examine decay factors inferred under different priors to guide exploration and interpretation of the AAV2 landscape. First, we found that the site-specific decay factors *δ_p_* inferred by the connectedness prior show substantial variation across positions, highlighting sites with little to no influence on the effects of other mutations (*δ*_575_ = 0.08, *δ*_561_ = 0.10) and sites with strong influence (*δ*_568_ = 0.37, *δ*_569_ = 0.52, Figure 3B). The Jenga prior exhibits even greater heterogeneity in the inferred decay factors, revealing that the identity of the mutated allele can matter more than the site itself. For example, while position 569 has the highest site-specific decay factor under the connectedness prior, the Jenga model shows that this effect is driven primarily by the Asn allele (*δ*^Asn^ = 0.43), with other substitutions at the same position showing minimal influence on predictability (*δ^a̸^*^=Asn^ *<* 0.08). To explore this further, we used the Gaussian process framework to compare posterior distributions of mutational effects in the wild-type background and in the presence of an Asn569Gln substitution (Figure S5A). In the wild-type background, mutations had widespread effects, but in the Asn569Gln background, these effects became largely neutral. This pattern reflects an essential role for Asn569 in capsid assembly, as all measured sequences lacking Asn at position 569 were non-functional (Figure 3C), with little apparent influence from genetic background (Figure S5B). Structurally, Asn569 forms hydrogen bonds with nearby residues, including 522 and 507 in AAV2 (Figure S5C), and these interactions are conserved across multiple AAV serotypes (Figure S5D, E). Even the conservative Asn569Gln substitution is predicted to introduce steric clashes that likely destabilize the capsid (Figure S5F).

Next, rather than focusing on a single position, we examined a region spanning positions 579–588, located in a solvent-exposed loop (Figure S6A), where the positively charged residues Arg and Lys exhibit consistently high decay rates (Figure 3B). In the wild-type background, mutational effect estimates reveal that Arg and Lys are generally deleterious throughout this region, whereas substitutions from positively charged to neutral or negatively charged residues, such as Glu and Asp, tend to be beneficial (Figure S6B). These patterns led us to hypothesize that the functional constraint in this region is charge-dependent. Supporting this hypothesis, mutations that reduce the overall charge, such as R588Q, are estimated to become neutral when introduced in an Asn587Glu background (Figure S6C) or even deleterious when combined with an additional Arg585Glu substitution (Figure S6D). To directly test this charge-balance hypothesis, we stratified the experimental data by the net charge in the 579-588 region. Consistent with our expectations, only sequences with a net charge between -4 and +2 could be functional, while those with excessive positive or negative charge were invariably non-functional. This suggests that an intermediate charge in this region is required, but not sufficient, for capsid assembly (Figure 3D).

We also examined the mutation-specific decay factors across all positions under the general product model (Figure S4). These values highlight the extensive variability and idiosyncrasy in amino acid exchangeability across different positions and biochemical contexts. Several positions exhibit mutation-specific decay factors that are incompatible with those expected under the Jenga prior, where mutations within groups are not expected to alter the effects of other mutations, whereas mutations across groups are. This pattern resembles position 54 in the GB1 dataset (Figure 1E). For instance, mutation-specific decay factors at position 576 exhibit two distinct groups: aromatic and non-aromatic residues (Figure 3E). Consistent with this hypothesis, the estimated mutational effects in the wild-type context were largely preserved in the presence of a Tyr576Phe substitution (Figure S5G), whereas other substitutions, like Tyr576Cys, systematically reduced the effects of other deleterious mutations (Figure S5H). Moreover, measured sequences without an aromatic at position 576 were generally non-functional (Figure 3F), thereby limiting the ability of other deleterious substitutions to further contribute to loss of function. This pattern is consistent with structural evidence: the wild-type Tyr576 occupies a hydrophobic pocket at the interface between two monomers, a location likely incompatible with non-aromatic residues (Figure S5I).

### Application to a genome-wide genotype-phenotype map

So far, we have applied our methods to genotype–phenotype maps for proteins and regulatory RNA sequences. Here, we extend our approach to a genome-wide genotype–phenotype map. We analyze a dataset derived from a large barcoded segregant library for haploid yeast [50]. Specifically, we modeled the relative fitness under lithium exposure for nearly 20,000 segregants with high-quality genotype data, focusing on the 83 previously identified quantitative trait loci (QTLs) spread across 15 chromosomes [50].

We fit the baseline additive, pairwise interaction, and geometric models to different proportions of the training data. All three models exhibit highly similar prediction performance across both low and high training data densities and converge to a test *R*^2^ of approximately 0.69 (Figure 4A). The failure of these epistatic models to substantially outperform the additive model suggests a lack of pervasive genetic interactions across genomic loci. We next fit connectedness regression, which in bi-allelic datasets such as this one is equivalent to both the Jenga and general product kernel regression models. Interestingly, we observed a consistent improvement in predictive performance relative to the baseline additive and epistatic models across all proportions of training data, achieving a test *R*^2^ of 0.76 on held-out data when including at least 50% of the data (Figure 4A). This result suggests that loci likely have heterogeneous epistatic contributions and differ in their influence on the predictability of mutations at other positions, highlighting the importance of accounting for this heterogeneity to achieve high predictive performance.

**Figure 4:**
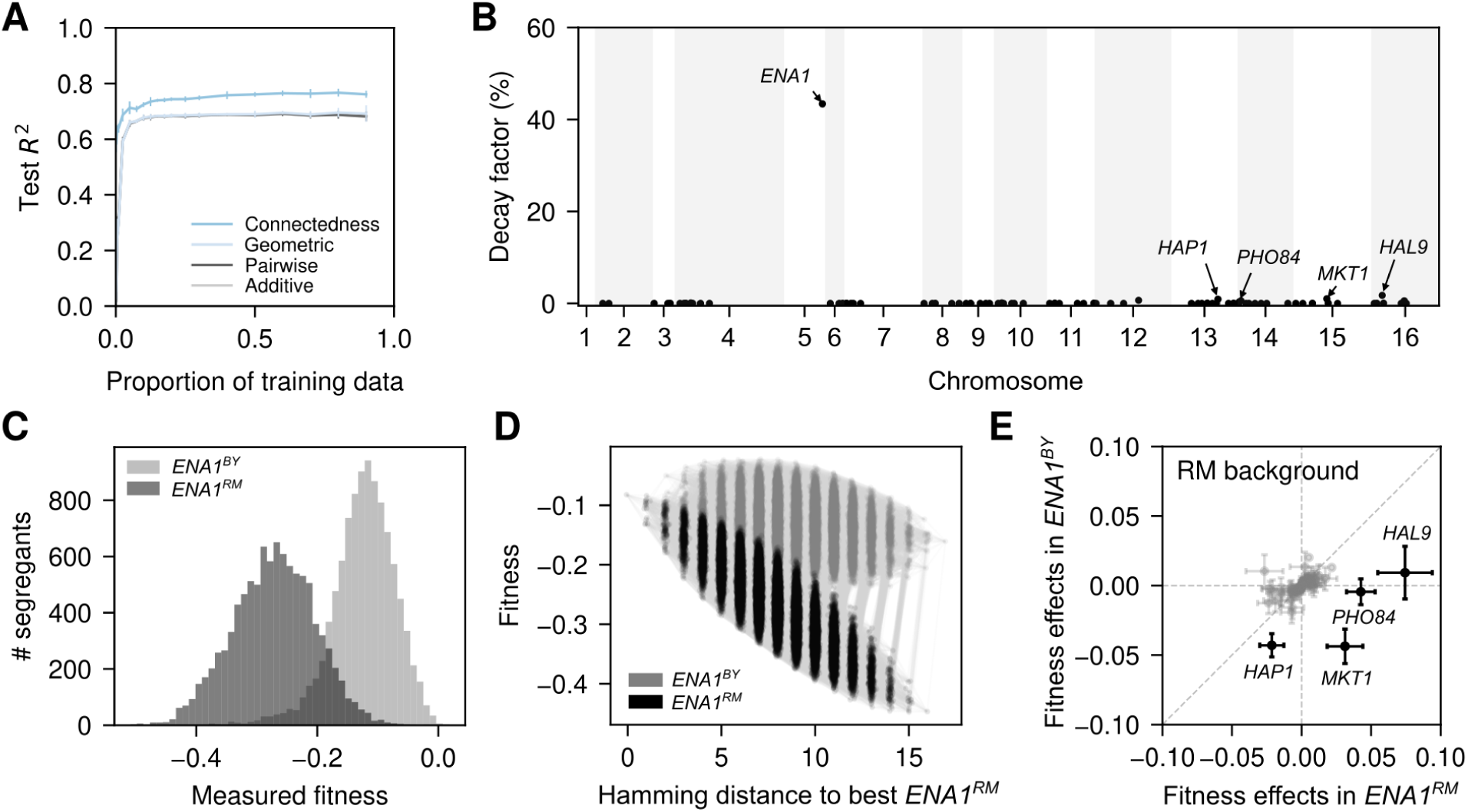
Connectedness regression applied to genome-wide genotype-phenotype data measuring relative fitness of *Saccharomyces cerevisiae* under lithium exposure. (A) Model predictive performance, measured by the *R*^2^ on held-out data, as a function of the fraction of data used for training. Error bars represent one standard deviation over three independent subsets of sequences used for training at each proportion. (B) Inferred decay factors for the 83 QTLs across the genome under the connectedness model. QTLs are named by the gene at which they are located or their closest gene in the reference genome S288C. (C) Fitness distribution of segregants stratified by their allele at the *ENA1* locus. (D) Representation of the complete genotype-phenotype map across the 16 loci with the highest inferred decay factors (mapped to the closest genes: *ENA1, HAL9, MKT1, PHO84, HAP1, HAL5, TAO3, BUL2, PTR2, NRT1, SUP45, DPH5, MLF3, SUS1, IRA2, VIP1*) and an additional pseudo-locus representing the genetic background across all other loci (RM vs. BY). The posterior mean fitnesses of the 2^17^ = 131,072 possible genotypes are plotted against their Hamming distances to the genotype with the highest predicted fitness when combined with the *ENA1* ^RM^ variant. Nodes represent genotypic combinations and are colored according to the allele at the *ENA1* locus. Edges connect genotypes separated by single point mutations. Values on the x-axis were jittered to facilitate visualization. (E) Comparison of the estimated mutational effects in the RM background in the presence of RM vs. BY alleles at the *ENA1* locus. Labeled QTLs highlighted in black correspond to loci with the largest decay factors after *ENA1*, as shown in (B). Error bars represent one standard deviation of the posterior distribution.

To further investigate the heterogeneous contribution of loci to the predictability of mutational effects, we examined locus-specific decay factors across the 83 focal QTLs (Figure 4B). Our analysis revealed that fitness variation is dominated by a single QTL located near the gene *ENA1* (*δ_ENA_*_1_ = 0.43), which encodes a Na^+^/Li^+^ ATPase pump involved in Na^+^ and Li^+^ efflux. As expected, this *ENA1* QTL also has a strong main effect, with genotypes carrying the beneficial BY or deleterious RM allele showing strongly shifted phenotypic distributions (Figure 4C).

To examine how the *ENA1* QTL reshapes the genotype–phenotype map, we computed the posterior distribution for the complete combinatorial landscape involving all possible genotypes at the 16 loci with the largest decay factors, with the background (BY or RM) encoded as an additional locus (Figure 4D). In contrast to the AAV2 example, where mutations at Asn569 caused complete loss of function and eliminated the effects of all other mutations, the deleterious *ENA1* ^RM^ allele modulates the effects of many mutations without uniformly suppressing them. Indeed, in this altered genetic context, previously neutral or deleterious variants can become beneficial, allowing combinations that achieve a relative fitness of *−*0.08, higher than the fitness of both the RM (*−*0.30) and BY (*−*0.18) wild-type strains.

To investigate how the *ENA1* QTL influences the effects of other QTLs in more detail, we computed the posterior distribution of the mutational effect for each of the other QTLs in the presence of *ENA1^RM^* or *ENA1^BY^* in an otherwise RM genetic background (Figure 4E). Whereas the effect of some QTLs with large decay factors, such as *HAP1*, remain deleterious in the presence of both *ENA1* alleles, we find 3 QTLs whose effects substantially change depending on the *ENA1* locus (Figures 4E and S7).

The first of these QTLs is near *HAL9*, a known regulator of *ENA1* expression [100]. We found that the *HAL9^BY^* allele is neutral in the *ENA1^BY^* background, but beneficial in the presence of the deleterious *ENA1^RM^* allele, suggesting that *HAL9^BY^* likely compensates for the deleterious *ENA1^RM^* via up-regulation of *ENA1*. Similarly, we found that for the second QTL, which mapped to PHO84, the BY allele is beneficial only in the *ENA1^RM^* background (Figure 4E). As *PHO84* is a high-affinity proton-phosphate symporter [101] and uses the proton gradient to import inorganic phosphate in the cell, it may influence the efficiency of a secondary detoxification mechanism involving the Na^+^, Li^+^-proton antiporter *NHA1*, which is known to become relevant in *ENA1* -deficient cells [100]. Finally, we found that the BY allele of a previously reported large-effect QTL near *MKT1* [50, 102] is detrimental in the presence of *ENA1^BY^*, but becomes beneficial when combined with the deleterious *ENA1^RM^* allele (Figure 4E). *MKT1* is a pleiotropic gene with known effects across multiple environments [50, 102], potentially by stabilizing specific mRNAs in a *PBP1* - and *PUF3* -dependent manner [103], although its interaction with genes involved in Li^+^ homeostasis, has not been previously reported to our knowledge. Importantly, the effects of these QTLs do not only depend on *ENA1*, but also vary across a wider range of genetic backgrounds (Figure S7). For instance, the beneficial effect of *HAL9^BY^* in the presence of *ENA1^RM^* is stronger in an RM rather than a BY genetic background across all other loci (Figure S7, upper left).

In summary, this yeast example demonstrates that our proposed models can infer genome-wide genotype-phenotype maps using data not only from engineered sequence libraries but also from high-throughput phenotyping of segregating variation in large populations. The high predictive performance of connectedness regression on held-out genotypes provides evidence of an important contribution of genetic interactions and coordinated epistasis to genome-scale genotype-phenotype maps, allowing us to detect and quantify novel genetic interactions and to exhaustively reconstruct small portions of even astronomically large genotype-phenotype maps (e.g., Figure 4D, showing a non-trivial sub-landscape consisting of 131,072 genotypes out of the 2^83^ *≈* 10^25^ possible genotypes in this system).

## Discussion

In this study, we introduced a family of models motivated by a simple yet informative aspect of epistasis: some mutations change the predictability of mutational effects more than others. Our methods provide a comprehensive framework for studying empirical sequence-function relationships by both allowing accurate phenotypic prediction for genotype-phenotype maps containing all orders of genetic interaction and providing biologically interpretable summaries of the structure of epistasis. The strong performance of these models across four empirical datasets, encompassing sequence-function relationships for proteins, RNAs, and complex traits, provides further evidence that heterogeneity across mutations in the degree to which they alter the effects of other mutations is likely a general feature of empirical sequence-function relationships, corresponding to genotype-phenotype maps that are rugged in some directions but smooth in others.

Our contribution also addresses a longstanding limitation of Gaussian process approaches: scalability to long sequences and large datasets. Classically, Gaussian processes could only be applied exactly to datasets containing fewer than low tens of thousands of observations [75], and while our previous methods exploited symmetries in biological sequence space to accommodate hundreds of thousands to low millions of observations[55, 58, 70, 74], they remained applicable only to short sequences. By leveraging recent advances in GPU acceleration[81–83], we can now overcome these previous constraints on sequence length, allowing us to provide software that can scale to longer protein sequences and genome-scale QTLs. Despite working in these astronomically large sequence spaces, our modeling framework maintains a high degree of interpretability because our hyperparameters have a definite genetic meaning in terms of how much each mutation reduces the predictability of other mutations. Moreover, by displaying these decay factors in heatmaps, we can at a glance identify the key alleles at each site most involved in epistatic interactions (e.g. Figure 3B) or subsets of amino acids that behave equivalently at a given site (e.g. Figure 3C, Figure S3), both of which can help guide the choice of mutations for follow-up experiments, and as we show, motivate biological hypotheses to explain the observed pattern of epistasis.

Our work also advances the ongoing process of unification and integration between theoretical and empirical approaches to fitness landscapes [5, 7, 8, 104]. In particular, an important advantage of Gaussian processes is that we can use well-characterized families of random fitness landscapes as priors for empirical data analysis [77, 85–88, 105]. By learning the hyperparameters for these priors via evidence maximization, our methods effectively identify the ensemble of theoretical landscapes whose patterns of epistasis are most similar to those in the observed data. At the same time, our proposal of the Jenga and general product kernel models provides an opportunity to analyze the dynamics of adaptive evolution and the accessibility of mutational paths under these more realistic forms of anisotropy [77]. Our work is also closely connected to the existing fitness landscape literature via our decay factors, which are conceptually related to the generalized *γ* statistics [28, 76] that measure the correlation in mutational effects across pairs of genetic backgrounds that differ by a specific mutation. We derive the mathematical relationship between the *γ* statistics and our decay factors in Supplement 1.3.

This work is also situated within a broader literature on Gaussian process approaches to modeling genotype-phenotype relationships [67–69, 106–111]. Previously, anisotropy has typically been introduced by incorporating external information, e.g. by building Gaussian processes over sequence embeddings [106, 107], by incorporating structural information [67], or by using information from sequence alignments [110]. By contrast, the models introduced here learn anisotropy directly from the data. In terms of expressiveness, our models are also similar in their motivations to deep neural networks that can act as general function approximators [34, 35, 46, 59–66], but Gaussian processes offer advantages in terms of interpretability and uncertainty quantification. One key advantage of DNNs is their ability to accommodate nonlinear transformations of model outputs, enabling them to capture global or nonspecific forms of epistasis [60, 61, 65, 112, 113]. In contrast, adding similar nonlinearities to Gaussian process models introduces non-Gaussian likelihoods requiring additional approximations, such as Laplace approximation [75] or variational inference [114–116]. Nonetheless, our new anisotropic Gaussian process models appear to consistently outperform global epistasis models except at very low training data densities. These new models also performed particularly well on the SMN1 dataset, which includes a strong global nonlinearity, and captured this nonlinear structure much better than our previous isotropic priors [55] While our models achieved excellent predictive performance, they also reveal several promising directions for future research. The connectedness, Jenga, and general product kernel priors by construction encode a form of “coordinated epistasis,” where the mutations with the largest effects also tend to be the most epistatic, a pattern consistent with many empirical observations [51, 117–120]. However, more work is needed to develop priors that capture not just how epistatic each mutation tends to be, but also provide more fine-grained summaries of which sites or mutations tend to interact with each other [121]. Another limitation is that while our priors allow anisotropy in the predictability of mutational effects, they are nonetheless homoskedastic, assigning uniform phenotypic variance across functionally distinct regions of sequence space. Developing heteroskedastic priors, where phenotypic variance varies across sequence space would better capture the expectation that functionally inert regions are less variable than those containing functionally active sequences. However, it is important to note that these are limitations on the inductive biases expressible through the prior, rather than limitations of the Bayesian regression models themselves. In particular, these models are capable of learning arbitrary genotype-phenotype maps, and any additional structure will still be reflected in the posterior, even if not directly encoded in the prior.

## Methods

### Data processing

**GB1 protein data** was processed as previously described [55, 58]. Briefly, we used the number of sequencing reads for each sequence *x* in the input sample (*c^input^*) and in the selected sample (*c^sel^*) reported in [84] to estimate the log-enrichment ratio relative to the wild-type sequence *y_x_* as a measure of the binding strength. Moreover, we estimated the error variance *σ*^2^ of this estimate [122]:

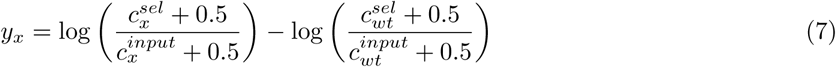

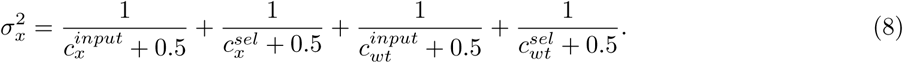

**SMN1** 5*^′^* **splice site data** was downloaded from https://github.com/davidmccandlish/vcregression, which used data from the original publication [42]. Briefly, assuming a log-normal distribution in 1-7 replicates, for each different 5*^′^* splice site sequence *x*, the bias corrected geometric mean of the enrichment ratio was used as an estimate of the median enrichment ratio when the enrichment ratio was strictly positive for all replicates. Otherwise, the median of enrichment scores was used to estimate the phenotype *y_x_*. Sequence-specific variance *σ*^2^ was estimated as indicated below, where *s*^2^ is the sample variance of the log-enrichment ratios if all replicates were strictly positive and were measured in at least two samples or the median of all *s*^2^ for sequences *x* with at least two replicate measurements:

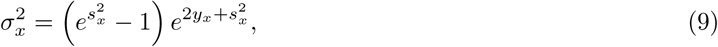

see [55] for more details.

**AAV2 capsid data** was downloaded from the ProteinGym database [123] and used as provided without external estimates of the experimental variance (the Gaussian process fit still included an experimental noise term with a learned uniform variance, see below).

**Yeast fitness data** was downloaded from the Dryad repository (https://doi.org/10.5061/dryad.1rn8pk0vd). This dataset provided estimates of the mean *y_x_* and variance *σ*^2^ of the relative fitness for each barcoded segregant *x*. We used the reported genotype probabilities *p_x,i_* for each position *i* across all *ℓ* loci to compute a segregant-level genome uncertainty *U_i_* as previously reported [50]

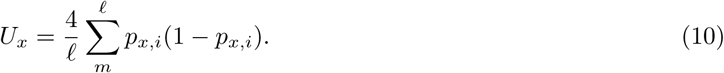

We kept only the 19833 (20%) segregants with lowest genome uncertainty for further analysis. We modeled the relative fitness in high Li^+^ concentration as a function of the 83 previously reported Quantitative Trait Loci (QTL) [50]. QTLs were labeled by associating them with the closest protein-coding gene within a 10kb window from the BY4741 genome annotation downloaded from the *Saccharomyces* Genome Database (SGD) [124].

### Global epistasis models

Global epistasis models for each data set were fit using MAVE-NN v1.0.2 [61] with an unregularized additive genotype-phenotype map, a monotonic global epistasis non-linearity, and Gaussian homoskedastic noise. Models were trained by minimizing the variational information with a batch size equal to the size of the dataset for 5000 epochs with a learning rate of 0.01 to ensure model convergence. All other parameters were set to their default values.

### Gaussian process models

Gaussian process models for the different datasets were fit using our newly developed library EpiK. EpiK implements the proposed prior distributions for performing Bayesian inference using GPyTorch [81] and KeOps [83] for GPU-accelerated inference of Gaussian process models. Specifically, we use a zero mean Gaussian process prior and a Gaussian likelihood with known experimental variance *σ*^2^ when available and fit an additional noise term *σ*^2^ for additional sequence-independent variance (see Supplement 1.1). Hyperparameters of each kernel were optimized by maximizing the Bayesian evidence using the Adam optimizer with a learning rate of 0.02 for 500 iterations. For accurate estimation of the evidence, we use a maximum number of 600 Lanczos vectors with 100 random vectors for trace estimation when approximating the log-determinant term (see Supplement 1.1). Site-specific product kernels (geometric, connectedness, Jenga and general product kernels) were initialized to have a prior variance equal to the empirical phenotypic variance and uniform decay factors across mutations and sites, which decay exponentially with the number of mutations down to 0.1 at the maximal distance between sequences. As the evidence is a non-convex function with potential local optima, to obtain the final estimates for the decay factors when fitting the complete dataset, we run 5 independent optimizations with different random initializations and kept the hyperparameter values that minimized the evidence along the 5 different optimizations. However, in practice, we see convergence to the same optima independent of the random initialization [75].

### Visualization of GB1 sequence-function map

We used the maximum a posteriori (MAP) estimate for the complete sequence-function map of protein GB1 inferred with the complete dataset under the Jenga model to generate a low-dimensional representation using gpmap-tools [74], in which squared distances optimally approximate the time to evolve between pairs of sequences under a model of molecular evolution where the population evolves under selection for higher log relative enrichment ratios. The different diffusion axes capture directions in which evolution is the slowest and typically separate qualitatively different sets of sequences with high phenotypic values that are separated by at least partial fitness valleys [58, 70, 93]. Selection strength was set such that the stationary distribution of the resulting Markov chain had a phenotypic mean that matches the wild-type phenotypic value.

## Acknowledgments

This work was supported by a Burroughs-Wellcome Careers at the Scientific Interface (CASI) award (SP), NIH grant R35 GM133613 (CMG, DMM), NIH grant R01 HG011787 (DMM), and NIH grant R35 GM154908 (JZ), as well as additional funding from the Simons Center for Quantitative Biology at CSHL (DMM) and the College of Liberal Arts and Sciences at the University of Florida (JZ).

## Data and code availability

The EpiK software package allows general Gaussian Process regression for sequence-function relationships under the kernels presented in this work. EpiK is freely available to the community at https://github.com/cmarti/epik under MIT license, with documentation available at https://epik.readthedocs.io. Data and code to reproduce the analysis and figures from this work are available at https://github.com/cmarti/epik_analyses.

## Notes

### Competing Interest Statement

The authors have declared no competing interest.

## References

1. S Wright. “The roles of mutation, inbreeding, crossbreeding and selection in evolution”. In: *Proceedings of the Sixth International Congress of Genetics*. 1932, pp. 356–366.

2. J Arjan G M De Visser and Joachim Krug. “Empirical fitness landscapes and the predictability of evolution”. In: *Nat. Rev. Genet.* 15.7 (2014), p. 480. doi: 10.1038/nrg3744.

[3] Patrick C Phillips. “Epistasis—the essential role of gene interactions in the structure and evolution of genetic systems”. In: Nat. Rev. Genet. 9.11 (2008), pp. 855–867. doi: 10.1038/nrg2452.

[4] Alexey S Kondrashov, Shamil Sunyaev, and Fyodor A Kondrashov. “Dobzhansky–Muller incompatibilities in protein evolution”. In: Proc. Natl. Acad. Sci. U.S.A. 99.23 (2002), pp. 14878–14883. doi: 10.1073/pnas.232565499.

[5] Daniel M Weinreich, Yinghong Lan, C Scott Wylie, and Robert B Heckendorn. “Should evolutionary geneticists worry about higher-order epistasis?” In: Curr. Opin. Genet. Dev. 23.6 (2013), pp. 700–707. doi: 10.1016/j.gde.2013.10.007.

[6] Zachary R Sailer and Michael J Harms. “High-order epistasis shapes evolutionary trajectories”. In: *PLoS Comput*. Biol. 13.5 (2017), e1005541. doi: 10.1371/journal.pcbi.1005541.

[7] Claudia Bank. “Epistasis and Adaptation on Fitness Landscapes”. en. In: Annual Review of Ecology, Evolution, and Systematics 53.Volume 53, 2022 (Nov. 2022), pp. 457–479. issn: 1543-592X, 1545-2069. doi: 10.1146/annurev-ecolsys-102320-112153.

[8] Milo S. Johnson, Gautam Reddy, and Michael M. Desai. “Epistasis and evolution: recent advances and an outlook for prediction”. en. In: BMC Biology 21.1 (May 2023), p. 120. issn: 1741-7007. doi: 10.1186/s12915-023-01585-3.

[9] Alief Moulana, Thomas Dupic, Angela M Phillips, Jeffrey Chang, Anne A Roffler, Allison J Greaney, Tyler N Starr, Jesse D Bloom, and Michael M Desai. “The landscape of antibody binding affinity in SARS-CoV-2 Omicron BA.1 evolution”. en. In: eLife 12 (Feb. 2023), e83442. issn: 2050-084X. doi: 10.7554/eLife.83442.

[10] Jason H. Moore and Scott M. Williams. “Epistasis and Its Implications for Personal Genetics”. English. In: The American Journal of Human Genetics 85.3 (Sept. 2009). Publisher: Elsevier, pp. 309–320. issn: 0002-9297, 1537-6605. doi: 10.1016/j.ajhg.2009.08.006. url: https://www.cell.com/ajhg/ abstract/S0002-9297(09)00349-8 (visited on 03/04/2025).

[11] Krishna Dasari, Jason A. Somarelli, Sudhir Kumar, and Jeffrey P. Townsend. “The somatic molecular evolution of cancer: Mutation, selection, and epistasis”. In: Progress in Biophysics and Molecular Biology. Cancer and Evolution 165 (Oct. 2021), pp. 56–65. issn: 0079-6107. doi: 10.1016/j.pbiomolbio.2021.08.003. url: https://www.sciencedirect.com/science/article/pii/S0079610721000961 (visited on 03/04/2025).

[12] Chase R Freschlin, Sarah A Fahlberg, and Philip A Romero. “Machine learning to navigate fitness landscapes for protein engineering”. In: Current Opinion in Biotechnology 75 (June 2022), p. 102713. issn: 0958-1669. doi: 10.1016/j.copbio.2022.102713. url: https://www.sciencedirect.com/ science/article/pii/S0958166922000465 (visited on 03/04/2025).

[13] Gloria Yang, Dave W Anderson, Florian Baier, Elias Dohmen, Nansook Hong, Paul D Carr, Shina Caroline Lynn Kamerlin, Colin J Jackson, Erich Bornberg-Bauer, and Nobuhiko Tokuriki. “Higher-order epistasis shapes the fitness landscape of a xenobiotic-degrading enzyme”. In: Nature Chemical Biology 15.11 (2019), pp. 1120–1128. doi: 10.1038/s41589-019-0386-3.

[14] Rosalie Lipsh-Sokolik and Sarel J. Fleishman. “Addressing epistasis in the design of protein function”. In: Proceedings of the National Academy of Sciences 121.34 (Aug. 2024). Publisher: Proceedings of the National Academy of Sciences, e2314999121. doi: 10.1073/pnas.2314999121. url: https://www.pnas.org/doi/10.1073/pnas.2314999121 (visited on 03/04/2025).

[15] Timothy B. Sackton and Daniel L. Hartl. “Genotypic Context and Epistasis in Individuals and Populations”. eng. In: Cell 166.2 (July 2016), pp. 279–287. issn: 1097-4172. doi: 10.1016/j.cell.2016.06.047.

[16] Gustavo De Los Campos, John M Hickey, Ricardo Pong-Wong, Hans D Daetwyler, and Mario P L Calus. “Whole-Genome Regression and Prediction Methods Applied to Plant and Animal Breeding”. en. In: Genetics 193.2 (Feb. 2013), pp. 327–345. issn: 1943-2631. doi: 10.1534/genetics.112.143313. url: https://academic.oup.com/genetics/article/193/2/327/6065337 (visited on 03/04/2025).

[17] Sebastian Soyk, Matthias Benoit, and Zachary B. Lippman. “New Horizons for Dissecting Epistasis in Crop Quantitative Trait Variation”. en. In: Annual Review of Genetics 54.Volume 54, 2020 (Nov. 2020). Publisher: Annual Reviews, pp. 287–307. issn: 0066-4197, 1545-2948. doi: 10.1146/annurev-genet-050720-122916. url: https://www.annualreviews.org/content/journals/10.1146/annurev-genet-050720-122916 (visited on 03/04/2025).

[18] Sangam L. Dwivedi, Pat Heslop-Harrison, Junrey Amas, Rodomiro Ortiz, and David Edwards. “Epistasis and pleiotropy-induced variation for plant breeding”. en. In: Plant Biotechnology Journal 22.10 (2024). _eprint: https://onlinelibrary.wiley.com/doi/pdf/10.1111/pbi.14405, pp. 2788–2807. issn: 1467-7652. doi: 10.1111/pbi.14405. url: https://onlinelibrary.wiley.com/doi/abs/10.1111/pbi.14405 (visited on 03/04/2025).

19. Justin B Kinney and David M McCandlish. “Massively Parallel Assays and Quantitative Sequence– Function Relationships”. In: *Annu. Rev. Genomics Hum. Genet*. 20 (2019).

[20] Douglas M Fowler, Carlos L Araya, Sarel J Fleishman, Elizabeth H Kellogg, Jason J Stephany, David Baker, and Stanley Fields. “High-resolution mapping of protein sequence-function relationships”. In: *Nat*. Methods 7.9 (2010), pp. 741–746. doi: 10.1038/nmeth.1492.

[21] Lea M Starita, Jonathan N Pruneda, Russell S Lo, Douglas M Fowler, Helen J Kim, Joseph B Hiatt, Jay Shendure, Peter S Brzovic, Stanley Fields, and Rachel E Klevit. “Activity-enhancing mutations in an E3 ubiquitin ligase identified by high-throughput mutagenesis”. In: Proc. Natl. Acad. Sci. U.S.A. 110.14 (2013), E1263–E1272. doi: 10.1073/pnas.1303309110.

[22] Daniel Melamed, David L Young, Caitlin E Gamble, Christina R Miller, and Stanley Fields. “Deep mutational scanning of an RRM domain of the *Saccharomyces cerevisiae* poly (A)-binding protein”. In: RNA 19.11 (2013), pp. 1537–1551. doi: 10.1261/rna.040709.113.

[23] C Anders Olson, Nicholas C Wu, and Ren Sun. “A comprehensive biophysical description of pairwise epistasis throughout an entire protein domain”. In: *Curr*. Biol. 24.22 (2014), pp. 2643–2651. doi: 10.1016/j.cub.2014.09.072.

[24] Michael B Doud, Orr Ashenberg, and Jesse D Bloom. “Site-specific amino acid preferences are mostly conserved in two closely related protein homologs”. In: Mol. Biol. Evol. 32.11 (2015), pp. 2944–2960. doi: 10.1093/molbev/msv167.

[25] Anna I Podgornaia and Michael T Laub. “Pervasive degeneracy and epistasis in a protein-protein interface”. In: Science 347.6222 (2015), pp. 673–677. doi: 10.1126/science.1257360.

[26] Karen S Sarkisyan, Dmitry A Bolotin, Margarita V Meer, Dinara R Usmanova, Alexander S Mishin, George V Sharonov, Dmitry N Ivankov, Nina G Bozhanova, Mikhail S Baranov, Onuralp Soylemez, et al. “Local fitness landscape of the green fluorescent protein”. In: Nature 533.7603 (2016), p. 397. doi: 10.1038/nature17995.

[27] Barrett Steinberg and Marc Ostermeier. “Shifting fitness and epistatic landscapes reflect trade-offs along an evolutionary pathway”. In: *J*. Mol. Biol. 428.13 (2016), pp. 2730–2743. doi: 10.1016/j.jmb. 2016.04.033.

[28] Claudia Bank, Sebastian Matuszewski, Ryan T Hietpas, and Jeffrey D Jensen. “On the (un)predictability of a large intragenic fitness landscape”. In: Proc. Natl. Acad. Sci. U.S.A. 113.49 (2016), pp. 14085–14090. doi: 10.1073/pnas.1612676113.

[29] Tyler N Starr, Lora K Picton, and Joseph W Thornton. “Alternative evolutionary histories in the sequence space of an ancient protein.” In: Nature 549.7672 (Sept. 2017), pp. 409–413. doi: 10.1038/ nature23902.

[30] Victoria O Pokusaeva, Dinara R Usmanova, Ekaterina V Putintseva, Lorena Espinar, Karen S Sarkisyan, Alexander S Mishin, Natalya S Bogatyreva, Dmitry N Ivankov, Arseniy V Akopyan, Sergey Ya Avvakumov, et al. “An experimental assay of the interactions of amino acids from orthologous sequences shaping a complex fitness landscape”. In: PLos Genet. 15.4 (2019), e1008079. doi: 10.1371/journal. pgen.1008079.

[31] Calin Plesa, Angus M Sidore, Nathan B Lubock, Di Zhang, and Sriram Kosuri. “Multiplexed gene synthesis in emulsions for exploring protein functional landscapes”. In: Science 359.6373 (2018), pp. 343– 347. doi: 10.1126/science.aao5167.

[32] Drew S Tack, Peter D Tonner, Abe Pressman, Nathan D Olson, Sasha F Levy, Eugenia F Romantseva, Nina Alperovich, Olga Vasilyeva, and David Ross. “The genotype-phenotype landscape of an allosteric protein”. In: Molecular systems biology 17.3 (2021), e10179. doi: 10.15252/msb.202010179.

[33] Tyler N Starr, Allison J Greaney, Sarah K Hilton, Daniel Ellis, Katharine HD Crawford, Adam S Dingens, Mary Jane Navarro, John E Bowen, M Alejandra Tortorici, Alexandra C Walls, et al. “Deep mutational scanning of SARS-CoV-2 receptor binding domain reveals constraints on folding and ACE2 binding”. In: Cell 182.5 (2020), pp. 1295–1310. doi: 10.1016/j.cell.2020.08.012.

[34] Louisa Gonzalez Somermeyer, Aubin Fleiss, Alexander S Mishin, Nina G Bozhanova, Anna A Igolkina, Jens Meiler, Maria-Elisenda Alaball Pujol, Ekaterina V Putintseva, Karen S Sarkisyan, and Fyodor A Kondrashov. “Heterogeneity of the GFP fitness landscape and data-driven protein design”. In: eLife 11 (2022). Ed. by Daniel J Kliebenstein and Naama Barkai, e75842. issn: 2050-084X. doi: 10.7554/eLife.75842.

[35] Drew H Bryant, Ali Bashir, Sam Sinai, Nina K Jain, Pierce J Ogden, Patrick F Riley, George M Church, Lucy J Colwell, and Eric D Kelsic. “Deep diversification of an AAV capsid protein by machine learning”. In: Nature Biotechnology 39.6 (2021), pp. 691–696. doi: 10.1038/s41587-020-00793-4.

[36] Andre J. Faure, Aina Martí-Aranda, Cristina Hidalgo-Carcedo, Antoni Beltran, Jörn M. Schmiedel, and Ben Lehner. “The genetic architecture of protein stability”. en. In: Nature 634.8035 (Oct. 2024). Publisher: Nature Publishing Group, pp. 995–1003. issn: 1476-4687. doi: 10.1038/s41586-024-07966-0. url: https://www.nature.com/articles/s41586-024-07966-0 (visited on 06/16/2025).

[37] Antoni Beltran, Xiang’er Jiang, Yue Shen, and Ben Lehner. “Site-saturation mutagenesis of 500 human protein domains”. en. In: Nature (Jan. 2025). Publisher: Nature Publishing Group, pp. 1–10. issn: 1476-4687. doi: 10.1038/s41586-024-08370-4.

[38] Justin B Kinney, Anand Murugan, Curtis G Callan, and Edward C Cox. “Using deep sequencing to characterize the biophysical mechanism of a transcriptional regulatory sequence”. In: Proc. Natl. Acad. Sci. U.S.A. 107.20 (2010), pp. 9158–9163. doi: 10.1073/pnas.1004290107.

[39] Alexander B. B. Rosenberg, Rupali P. P. Patwardhan, Jay Shendure, and Georg Seelig. “Learning the Sequence Determinants of Alternative Splicing from Millions of Random Sequences”. In: Cell 163.3 (Oct. 2015). Publisher: Cell Press, pp. 698–711. issn: 10974172. doi: 10.1016/j.cell.2015.09.054.

[40] Michal Rabani, Lindsey Pieper, Guo-Liang Chew, and Alexander F. Schier. “A Massively Parallel Reporter Assay of 3 UTR Sequences Identifies In Vivo Rules for mRNA Degradation”. English. In: Molecular Cell 68.6 (Dec. 2017). Publisher: Elsevier, 1083–1094.e5. issn: 1097-2765. doi: 10.1016/j. molcel.2017.11.014.

[41] Sergey A. Evfratov, Ilya A. Osterman, Ekaterina S. Komarova, Alexandra M. Pogorelskaya, Maria P. Rubtsova, Timofei S. Zatsepin, Tatiana A. Semashko, Elena S. Kostryukova, Andrey A. Mironov, Evgeny Burnaev, Ekaterina Krymova, Mikhail S. Gelfand, Vadim M. Govorun, Alexey A. Bogdanov, Petr V. Sergiev, and Olga A. Dontsova. “Application of sorting and next generation sequencing to study 5’-UTR influence on translation efficiency in Escherichia coli”. In: Nucleic Acids Research 45.6 (2017), pp. 3487–3502. issn: 13624962. doi: 10.1093/nar/gkw1141.

[42] Mandy S Wong, Justin B Kinney, and Adrian R Krainer. “Quantitative Activity Profile and Context Dependence of All Human 5’ Splice Sites”. In: *Mol*. Cell (2018). doi: 10.1016/j.molcel.2018.07.033.

[43] Pablo Baeza-Centurion, Belén Miñana, Jörn M. Schmiedel, Juan Valcárcel, and Ben Lehner. “Combinatorial Genetics Reveals a Scaling Law for the Effects of Mutations on Splicing”. In: Cell 176.3 (Jan. 2019). Publisher: Cell Press, 549–563.e23. issn: 10974172. doi: 10.1016/j.cell.2018.12.010. url: https://www.sciencedirect.com/science/article/pii/S0092867418316246?via%3Dihub (visited on 03/18/2019).

[44] Carl G. de Boer, Eeshit Dhaval Vaishnav, Ronen Sadeh, Esteban Luis Abeyta, Nir Friedman, and Aviv Regev. “Deciphering eukaryotic gene-regulatory logic with 100 million random promoters”. en. In: Nature Biotechnology 38.1 (Jan. 2020). Publisher: Nature Publishing Group, pp. 56–65. issn: 1546-1696. doi: 10.1038/s41587-019-0315-8. url: https://www.nature.com/articles/s41587-019-0315-8 (visited on 06/16/2025).

[45] Syue-Ting Kuo, Ruey-Lin Jahn, Yuan-Ju Cheng, Yi-Lan Chen, Yun-Ju Lee, Florian Hollfelder, Jin-Der Wen, and Hsin-Hung David Chou. “Global fitness landscapes of the Shine-Dalgarno sequence”. In: Genome Research 30.5 (2020), pp. 711–723. doi: 10.1101/gr.260182.119.

[46] Eeshit Dhaval Vaishnav, Carl G. de Boer, Jennifer Molinet, Moran Yassour, Lin Fan, Xian Adiconis, Dawn A. Thompson, Joshua Z. Levin, Francisco A. Cubillos, and Aviv Regev. “The evolution, evolvability and engineering of gene regulatory DNA”. In: Nature (2022). Publisher: Springer US. issn: 0028-0836. doi: 10.1038/s41586-022-04506-6.

[47] Susan E. Liao, Mukund Sudarshan, and Oded Regev. “Deciphering RNA splicing logic with interpretable machine learning”. en. In: Proceedings of the National Academy of Sciences 120.41 (Oct. 2023), e2221165120. issn: 0027-8424, 1091-6490. doi: 10.1073/pnas.2221165120.

[48] Vikram Agarwal, Fumitaka Inoue, Max Schubach, Dmitry Penzar, Beth K. Martin, Pyaree Mohan Dash, Pia Keukeleire, Zicong Zhang, Ajuni Sohota, Jingjing Zhao, Ilias Georgakopoulos-Soares, William S. Noble, Galip Gürkan Yardımcı, Ivan V. Kulakovskiy, Martin Kircher, Jay Shendure, and Nadav Ahituv. “Massively parallel characterization of transcriptional regulatory elements”. en. In: Nature (Jan. 2025). Publisher: Nature Publishing Group, pp. 1–10. issn: 1476-4687. doi: 10.1038/s41586-024-08430-9.

[49] Christopher W Bakerlee, Alex N Nguyen, Yekaterina Shulgina, Jose I Rojas Echenique, and Michael M Desai. “Idiosyncratic epistasis leads to global fitness–correlated trends”. en. In: *Science* (2022).

50. Alex N Nguyen Ba, Katherine R Lawrence, Artur Rego-Costa, Shreyas Gopalakrishnan, Daniel Temko, Franziska Michor, and Michael M Desai. “Barcoded bulk QTL mapping reveals highly polygenic and epistatic architecture of complex traits in yeast”. en. In: *eLife* 11 (Feb. 2022), e73983. issn: 2050-084X.

51. Takeshi Matsui, Martin N Mullis, Kevin R Roy, Joseph J Hale, Rachel Schell, Sasha F Levy, and Ian M Ehrenreich. “The interplay of additivity, dominance, and epistasis on fitness in a diploid yeast cross”. In: *Nature Communications* 13.1 (2022), p. 1463.

52. Arnaud N’Guessan, Wen Yuan Tong, Hamed Heydari, and Alex N. Nguyen Ba. “Refining the resolution of the yeast genotype-phenotype map using single-cell RNA-sequencing”. en. In: *eLife* 13 (May 2025). Publisher: eLife Sciences Publications Limited.

53. Dmitry A Kondrashov and Fyodor A Kondrashov. “Topological features of rugged fitness landscapes in sequence space”. In: *Trends Genet*. 31.1 (2015), pp. 24–33.

54. Júlia Domingo, Pablo Baeza-Centurion, and Ben Lehner. “The Causes and Consequences of Genetic Interactions (Epistasis)”. In: *Annu. Rev. Genomics Hum. Genet*. 20 (2019).

55. Juannan Zhou, Mandy S Wong, Wei-Chia Chen, Adrian R Krainer, Justin B Kinney, and David M McCandlish. “Higher-order epistasis and phenotypic prediction”. In: *Proceedings of the National Academy of Sciences* 119.39 (2022), e2204233119.

56. Yeonwoo Park, Brian PH Metzger, and Joseph W Thornton. “Epistatic drift causes gradual decay of predictability in protein evolution”. In: *Science* 376.6595 (2022), pp. 823–830.

57. R A Fisher. “The Correlation Between Relatives on the Supposition of Mendelian Inheritance”. In: *Trans. R. Soc. Edinburgh* 52.02 (1918), pp. 399–433.

58. Juannan Zhou and David M McCandlish. “Minimum epistasis interpolation for sequence-function relationships”. In: *Nature Communications* 11.1 (2020), pp. 1–14.

59. Amirali Aghazadeh, Hunter Nisonoff, Orhan Ocal, David H Brookes, Yijie Huang, O Ozan Koyluoglu, Jennifer Listgarten, and Kannan Ramchandran. “Epistatic Net allows the sparse spectral regularization of deep neural networks for inferring fitness functions”. In: *Nature Communications* 12.1 (2021), pp. 1– 10.

60. Karen S Sarkisyan, Dmitry A Bolotin, Margarita V Meer, Dinara R Usmanova, Alexander S Mishin, George V Sharonov, Dmitry N Ivankov, Nina G Bozhanova, Mikhail S Baranov, Onuralp Soylemez, et al. “Local fitness landscape of the green fluorescent protein”. In: *Nature* 533.7603 (2016), pp. 397–401.

61. Ammar Tareen, Mahdi Kooshkbaghi, Anna Posfai, William T Ireland, David M McCandlish, and Justin B Kinney. “MAVE-NN: learning genotype-phenotype maps from multiplex assays of variant effect”. In: *Genome biology* 23.1 (2022), p. 98.

62. Sam Gelman, Sarah A. Fahlberg, Pete Heinzelman, Philip A. Romero, and Anthony Gitter. “Neural networks to learn protein sequence–function relationships from deep mutational scanning data”. en. In: *Proceedings of the National Academy of Sciences* 118.48 (Nov. 2021), e2104878118. issn: 0027-8424, 1091-6490.

[63] Bernardo P. de Almeida, Franziska Reiter, Michaela Pagani, and Alexander Stark. “DeepSTARR predicts enhancer activity from DNA sequence and enables the de novo design of synthetic enhancers”. en. In: Nature Genetics 54.5 (May 2022). Publisher: Nature Publishing Group, pp. 613–624. issn: 1546-1718. doi: 10.1038/s41588-022-01048-5. url: https://www.nature.com/articles/s41588-022-01048-5 (visited on 06/16/2025).

64. Chase R. Freschlin, Sarah A. Fahlberg, Pete Heinzelman, and Philip A. Romero. “Neural network extrapolation to distant regions of the protein fitness landscape”. en. In: *Nature Communications* 15.1 (July 2024), p. 6405. issn: 2041-1723.

[65] Palash Sethi and Juannan Zhou. “Importance of higher-order epistasis in large protein sequence-function relationships”. In: *bioRxiv* (2024).

[66] Mike Thompson, Mariano Martín, Trinidad Sanmartín Olmo, Chandana Rajesh, Peter K Koo, Benedetta Bolognesi, and Ben Lehner. “Massive experimental quantification allows interpretable deep learning of protein aggregation”. en. In: *Science Advances* (2025).

67. Philip A Romero, Andreas Krause, and Frances H Arnold. “Navigating the protein fitness landscape with Gaussian processes”. In: *Proc. Natl. Acad. Sci. U.S.A*. 110.3 (2013), E193–E201.

68. Daniel Gianola and Johannes BCHM Van Kaam. “Reproducing kernel Hilbert spaces regression methods for genomic assisted prediction of quantitative traits”. In: *Genetics* 178.4 (2008), pp. 2289–2303.

69. Gota Morota, Masanori Koyama, Guilherme J M Rosa, Kent A Weigel, and Daniel Gianola. “Predicting complex traits using a diffusion kernel on genetic markers with an application to dairy cattle and wheat data”. In: *Genetics Selection Evolution* 45 (2013), pp. 1–15.

70. Wei-Chia Chen, Juannan Zhou, Jason M Sheltzer, Justin B Kinney, and David M McCandlish. “Field-theoretic density estimation for biological sequence space with applications to 5 splice site diversity and aneuploidy in cancer”. In: *Proceedings of the National Academy of Sciences* 118.40 (2021).

71. Wei-Chia Chen, Juannan Zhou, and David M McCandlish. “Density estimation for ordinal biological sequences and its applications”. In: *Physical Review E* 110.4 (2024), p. 044408.

[72] Jacob T. Rapp, Bennett J. Bremer, and Philip A. Romero. “Self-driving laboratories to autonomously navigate the protein fitness landscape”. en. In: Nature Chemical Engineering 1.1 (Jan. 2024). Publisher: Nature Publishing Group, pp. 97–107. issn: 2948-1198. doi: 10.1038/s44286-023-00002-4. url: https://www.nature.com/articles/s44286-023-00002-4 (visited on 06/17/2025).

73. Samantha Petti, Carlos Martí-Gómez, Justin B Kinney, Juannan Zhou, and David M McCandlish. “On learning functions over biological sequence space: relating Gaussian process priors, regularization, and gauge fixing”. In: *arXiv preprint arXiv:2504.19034* (2025).

74. Carlos Martí-Gómez, Juannan Zhou, Wei-Chia Chen, Justin B. Kinney, and David M. McCandlish. “Inference and visualization of complex genotype-phenotype maps with *gpmap-tools*”. en. In: *bioRxiv* (Mar. 2025).

75. Carl Edward Rasmussen and Christopher K I Williams. *Gaussian processes for machine learning*. MIT Press, 2006.

76. Luca Ferretti, Benjamin Schmiegelt, Daniel Weinreich, Atsushi Yamauchi, Yutaka Kobayashi, Fumio Tajima, and Guillaume Achaz. “Measuring epistasis in fitness landscapes: The correlation of fitness effects of mutations”. In: *J. Theor. Biol*. 396 (2016), pp. 132–143.

77. Gautam Reddy and Michael M Desai. “Global epistasis emerges from a generic model of a complex trait”. In: *Elife* 10 (2021), e64740.

78. Daniel M Weinreich, Richard A Watson, and Lin Chao. “Perspective: sign epistasis and genetic costraint on evolutionary trajectories”. In: *Evolution* 59.6 (2005), pp. 1165–1174.

79. Daniel J Kvitek and Gavin Sherlock. “Reciprocal sign epistasis between frequently experimentally evolved adaptive mutations causes a rugged fitness landscape”. In: *PLoS genetics* 7.4 (2011), e1002056.

[80] John FC Kingman. “A simple model for the balance between selection and mutation”. In: Journal of Applied Probability 15.1 (1978), pp. 1–12.

81. Jacob Gardner, Geoff Pleiss, Kilian Q Weinberger, David Bindel, and Andrew G Wilson. “Gpytorch: Blackbox matrix-matrix gaussian process inference with gpu acceleration”. In: *Advances in neural information processing systems* 31 (2018).

[82] Ke Wang, Geoff Pleiss, Jacob Gardner, Stephen Tyree, Kilian Q Weinberger, and Andrew Gordon Wilson. “Exact Gaussian Processes on a Million Data Points”. en. In: *NeurIPS* (2019).

[83] Benjamin Charlier, Jean Feydy, Joan Alexis Glaunes, François-David Collin, and Ghislain Durif. “Kernel operations on the GPU, with autodiff, without memory overflows”. In: Journal of Machine Learning Research 22.74 (2021), pp. 1–6.

[84] Nicholas C. Wu, C. Anders Olson, and Ren Sun. “High-throughput identification of protein mutant stability computed from a double mutant fitness landscape”. In: Protein Sci. 25.2 (2016), pp. 530–539. issn: 1469896X. doi: 10.1002/pro.2840.

[85] Peter F Stadler, Robert Happel, et al. “Canonical approximation of landscapes”. In: *Santa Fe Institute Preprint* (1994), pp. 94–09.

86. Peter F Stadler and Robert Happel. “Random field models for fitness landscapes”. In: *J. Math. Biol*. 38.5 (1999), pp. 435–478.

[87] Johannes Neidhart, Ivan G Szendro, and Joachim Krug. “Exact results for amplitude spectra of fitness landscapes”. In: Journal of Theoretical Biology 332 (2013), pp. 218–227.

88. Atish Agarwala and Daniel S Fisher. “Adaptive walks on high-dimensional fitness landscapes and seascapes with distance-dependent statistics”. In: *Theoretical Population Biology* 130 (2019), pp. 13–49.

89. Tinghua Wang, Dongyan Zhao, and Shengfeng Tian. “An overview of kernel alignment and its applications”. In: *Artificial Intelligence Review* 43 (2015), pp. 179–192.

90. Risi Imre Kondor and John D. Lafferty. “Diffusion Kernels on Graphs and Other Discrete Input Spaces”. In: *Proceedings of the Nineteenth International Conference on Machine Learning*. ICML ’02. San Francisco, CA, USA: Morgan Kaufmann Publishers Inc., July 2002, pp. 315–322. isbn: 978-1-55860-873-3. (Visited on 08/05/2025).

[91] Radford M. Neal. *Bayesian Learning for Neural Networks*. Ed. by P. Bickel, P. Diggle, S. Fienberg, K. Krickeberg, I. Olkin, N. Wermuth, and S. Zeger. Vol. 118. Lecture Notes in Statistics. New York, NY: Springer, 1996. doi: 10.1007/978-1-4612-0745-0. url: http://link.springer.com/10.1007/978-1-4612-0745-0 (visited on 06/16/2025).

[92] Daniel Lewandowski, Dorota Kurowicka, and Harry Joe. “Generating random correlation matrices based on vines and extended onion method”. In: Journal of Multivariate Analysis 100.9 (Oct. 2009), pp. 1989– 2001. issn: 0047-259X. doi: 10.1016/j.jmva.2009.04.008. url: https://www.sciencedirect.com/ science/article/pii/S0047259X09000876 (visited on 12/16/2024).

93. David M McCandlish. “Visualizing fitness landscapes”. In: *Evolution* 65.6 (2011), pp. 1544–1558.

94. Jakub Otwinowski, David Martin McCandlish, and Joshua B Plotkin. “Inferring the shape of global epistasis”. In: *Proceedings of the National Academy of Sciences* 115.32 (2018), E7550–E7558.

95. Zachary R Sailer and Michael J Harms. “Detecting high-order epistasis in nonlinear genotype-phenotype maps”. In: *Genetics* 205.3 (2017), pp. 1079–1088.

96. Anna Posfai, Juannan Zhou, David M McCandlish, and Justin B Kinney. “Gauge fixing for sequence-function relationships”. In: *PLoS Computational Biology* 21.3 (2025), e1012818.

97. Tyler N Starr and Joseph W Thornton. “Epistasis in protein evolution”. In: *Protein Sci*. 25.7 (2016), pp. 1204–1218.

[98] Yuma Ishigami, Mandy S. Wong, Carlos Martí-Gómez, Andalus Ayaz, Mahdi Kooshkbaghi, Sonya M. Hanson, David M. McCandlish, Adrian R. Krainer, and Justin B. Kinney. “Specificity, synergy, and mechanisms of splice-modifying drugs”. en. In: Nature Communications 15.1 (Feb. 2024), p. 1880. issn: 2041-1723. doi: 10.1038/s41467-024-46090-5. url: https://www.nature.com/articles/s41467-024-46090-5 (visited on 07/16/2024).

99. Sam Sinai, Nina Jain, George M Church, and Eric D Kelsic. “Generative AAV capsid diversification by latent interpolation”. en. In: *bioRxiv* (Apr. 2021). doi: 10.1101/2021.04.16.440236. (Visited on 02/02/2024).

100. Amparo Ruiz and Joaquín Ariño. “Function and Regulation of the Saccharomyces cerevisiae ENA Sodium ATPase System”. en. In: *Eukaryotic Cell* 6.12 (Oct. 2007), p. 2175.

101. Elja Eskes, Marie-Anne Deprez, Tobias Wilms, and Joris Winderickx. “pH homeostasis in yeast; the phosphate perspective”. en. In: *Current Genetics* 64.1 (Feb. 2018), pp. 155–161. issn: 1432-0983.

[102] Adam M. Deutschbauer and Ronald W. Davis. “Quantitative trait loci mapped to single-nucleotide resolution in yeast”. en. In: Nature Genetics 37.12 (Dec. 2005). Publisher: Nature Publishing Group, pp. 1333–1340. issn: 1546-1718. doi: 10.1038/ng1674.

103. Koppisetty Viswa Chaithanya and Himanshu Sinha. “MKT1 alleles regulate stress responses through posttranscriptional modulation of Puf3 targets in budding yeast”. en. In: *Yeast* 40.12 (2023). _eprint: https://onlinelibrary.wiley.com/doi/pdf/10.1002/yea.3908, pp. 616–627. issn: 1097-0061.

[104] Ivan G Szendro, Martijn F Schenk, Jasper Franke, Joachim Krug, and J Arjan G M de Visser. “Quantitative analyses of empirical fitness landscapes”. In: J Stat Mech Theory Exp 2013.01 (2013), P01005. doi: 10.1088/1742-5468/2013/01/P01005.

105. Johannes Neidhart, Ivan G Szendro, and Joachim Krug. “Adaptation in Tunably Rugged Fitness Landscapes: The Rough Mount Fuji Model”. en. In: *Genetics* 198.2 (Oct. 2014), pp. 699–721. issn: 1943-2631.

106. Kevin K Yang, Zachary Wu, and Frances H Arnold. “Machine-learning-guided directed evolution for protein engineering”. In: *Nat. Methods* 16.8 (2019), pp. 687–694.

107. Peter Mørch Groth, Mads Herbert Kerrn, Lars Olsen, Jesper Salomon, and Wouter Boomsma. “Kermut: Composite kernel regression for protein variant effects”. en. In: *arXiv* (May 2024).

108. Christina Leslie, Eleazar Eskin, and William Stafford Noble. “THE SPECTRUM KERNEL: A STRING KERNEL FOR SVM PROTEIN CLASSIFICATION”. en. In: *Biocomputing* 2002. Kauai, Hawaii, USA: WORLD SCIENTIFIC, Dec. 2001, pp. 564–575.

109. Christina S. Leslie, Eleazar Eskin, Adiel Cohen, Jason Weston, and William Stafford Noble. “Mismatch string kernels for discriminative protein classification”. In: *Bioinformatics* 20.4 (Mar. 2004), pp. 467–476. issn: 1367-4803.

110. David Haussler. “Convolution Kernels on Discrete Structures”. In: (1999).

[111] Alan Nawzad Amin, Eli Nathan Weinstein, and Debora Susan Marks. *Biological Sequence Kernels with Guaranteed Flexibility*. en. arXiv:2304.03775 [cs, q-bio, stat]. Apr. 2023.

112. Louisa Gonzalez Somermeyer, Aubin Fleiss, Alexander S Mishin, Nina G Bozhanova, Anna A Igolkina, Jens Meiler, Maria-Elisenda Alaball Pujol, Ekaterina V Putintseva, Karen S Sarkisyan, and Fyodor A Kondrashov. “Heterogeneity of the GFP fitness landscape and data-driven protein design”. In: *Elife* 11 (2022), e75842.

113. Andre J. Faure and Ben Lehner. “MoCHI: neural networks to fit interpretable models and quantify energies, energetic couplings, epistasis, and allostery from deep mutational scanning data”. In: *Genome Biology* 25.1 (Dec. 2024), p. 303. issn: 1474-760X.

114. James Hensman, Alex Matthews, and Zoubin Ghahramani. “Scalable Variational Gaussian Process Classification”. en. In: *arXiv.org* (Nov. 2014).

[115] Alp Kucukelbir, Dustin Tran, Rajesh Ranganath, Andrew Gelman, and David M. Blei. “Automatic Differentiation Variational Inference”. In: *arXiv* (2016), pp. 1–38. issn: 15337928.

116. Peter D Tonner, Abe Pressman, and David Ross. “Interpretable modeling of genotype–phenotype landscapes with state-of-the-art predictive power”. In: *Proceedings of the National Academy of Sciences* 119.26 (2022), e2114021119.

117. Michael Costanzo, Anastasia Baryshnikova, Jeremy Bellay, Yungil Kim, Eric D. Spear, Carolyn S. Sevier, Huiming Ding, Judice L.Y. Koh, Kiana Toufighi, Sara Mostafavi, Jeany Prinz, Robert P. St. Onge, Benjamin VanderSluis, Taras Makhnevych, Franco J. Vizeacoumar, Solmaz Alizadeh, Sondra Bahr, Renee L. Brost, Yiqun Chen, Murat Cokol, Raamesh Deshpande, Zhijian Li, Zhen-Yuan Lin, Wendy Liang, Michaela Marback, Jadine Paw, Bryan-Joseph San Luis, Ermira Shuteriqi, Amy Hin Yan Tong, Nydia van Dyk, Iain M. Wallace, Joseph A. Whitney, Matthew T. Weirauch, Guoqing Zhong, Hongwei Zhu, Walid A. Houry, Michael Brudno, Sasan Ragibizadeh, Balázs Papp, Csaba Pál, Frederick P. Roth, Guri Giaever, Corey Nislow, Olga G. Troyanskaya, Howard Bussey, Gary D. Bader, Anne-Claude Gingras, Quaid D. Morris, Philip M. Kim, Chris A. Kaiser, Chad L. Myers, Brenda J. Andrews, and Charles Boone. “The Genetic Landscape of a Cell”. In: *Science* 327.5964 (Jan. 2010), pp. 425–431.

118. Joshua S. Bloom, Iulia Kotenko, Meru J. Sadhu, Sebastian Treusch, Frank W. Albert, and Leonid Kruglyak. “Genetic interactions contribute less than additive effects to quantitative trait variation in yeast”. en. In: *Nature Communications* 6.1 (Nov. 2015). Publisher: Nature Publishing Group, p. 8712. issn: 2041-1723.

119. Brooke Sheppard, Nadav Rappoport, Po-Ru Loh, Stephan J Sanders, Noah Zaitlen, and Andy Dahl. “A model and test for coordinated polygenic epistasis in complex traits”. In: *Proceedings of the National Academy of Sciences* 118.15 (2021), e1922305118.

[120] David Tang, Jerome Freudenberg, and Andy Dahl. “Factorizing polygenic epistasis improves prediction and uncovers biological pathways in complex traits”. In: The American Journal of Human Genetics 110.11 (2023), pp. 1875–1887.

[121] Sungmin Hwang, Benjamin Schmiegelt, Luca Ferretti, and Joachim Krug. “Universality Classes of Interaction Structures for NK Fitness Landscapes”. In: Journal of Statistical Physics 172.1 (2018). arXiv: 1708.06556 Publisher: Springer US, pp. 226–278. issn: 00224715.

122. Alan F Rubin, Hannah Gelman, Nathan Lucas, Sandra M Bajjalieh, Anthony T Papenfuss, Terence P Speed, and Douglas M Fowler. “A statistical framework for analyzing deep mutational scanning data”. In: *Genome Biol*. 18.1 (2017), p. 150.

123. Pascal Notin, Aaron W. Kollasch, Daniel Ritter, Lood Van Niekerk, Steffanie Paul, Hansen Spinner, Nathan Rollins, Ada Shaw, Ruben Weitzman, Jonathan Frazer, Mafalda Dias, Dinko Franceschi, Rose Orenbuch, Yarin Gal, and Debora S. Marks. “ProteinGym: Large-Scale Benchmarks for Protein Design and Fitness Prediction”. en. In: *bioRxiv* (Dec. 2023). doi: 10.1101/2023.12.07. 570727.

[124] Stacia R. Engel, Edith D. Wong, Robert S. Nash, Suzi Aleksander, Micheal Alexander, Eric Douglass, Kalpana Karra, Stuart R. Miyasato, Matt Simison, Marek S. Skrzypek, Shuai Weng, and J. Michael Cherry. “New data and collaborations at the Saccharomyces Genome Database: updated reference genome, alleles, and the Alliance of Genome Resources”. eng. In: Genetics 220.4 (Apr. 2022), iyab224. issn: 1943-2631. doi: 10.1093/genetics/iyab224.

