## Supplementary Information for "Learning sequence-function relationships with scalable, interpretable Gaussian processes"

### 1 Supplementary information

#### 1.1 Introduction to Gaussian Process regression

In this section, we present a brief overview of the Gaussian process regression framework in the context of modeling sequence-function relationships.

Gaussian processes (GP) provide a highly flexible class of non-parametric methods appropriate for modeling sequence-function data [1–3]. To construct a GP model, we begin by specifying a prior distribution over the space of all functions, such that the probability of sampling a function $f$ is given by a zero mean Gaussian distribution

$$f \sim \mathcal{N}(0, K), \tag{1}$$

where $K$ is the covariance matrix. Specifically, entries of $K$ are given by the so-called kernel function which measures the similarity between pairs of sequences $x, x'$ in our prior, such that $K_{x,x'} = k(x, x')$.

Given phenotypic measurements for a set of sequences $B$ arranged on a vector $y \in \mathbb{R}^{|B|}$, we predict the phenotype for any genotype by calculating its posterior distribution given the data, which again is Gaussian, with mean

$$\mathbb{E}[f(x)|\, y] = k_x^T (K_{BB} + E)^{-1} y, \tag{2}$$

where $K_{BB}$ is the submatrix of $K$ indexed by sequences in $B$ and $E$ is a diagonal matrix describing the experimental noise. $E_{x,x} = \sigma_x^2 + \sigma_n^2$, where $\sigma_x^2$ is the previously known variance of the measurement at sequence $x$ and $\sigma_n^2$ is a hyperparameter representing an additional noise term. When independent genotype-level noise estimates $\sigma_x^2$ are unavailable, we set $\sigma_x^2 = 0$ such that the hyperparameter $\sigma_n^2$ fully represents homoskedastic experimental noise. $k_x^T$ is the row vector containing kernel entries between the focal genotype $x$ and all genotypes in the data $B$. One major advantage of GP regression models over parametric methods such as neural networks is that GP regression provides a principled method for estimating the prediction uncertainty through direct calculation of the posterior covariance matrix

$$\mathrm{Var}[f(x)|\, y] = k(x, x) - k_x^T (K_{BB} + E)^{-1} k_x. \tag{3}$$

The critical step in implementing a GP model for sequence-function relationships is to specify an appropriate prior distribution (equivalently, a kernel function) that can adequately capture the large scale statistical characteristics of the data. For example, in our previous method empirical variance component regression (VC regression [2]), the kernel function is

$$k(x, x') = \sum_{k=1}^{\ell} \lambda_k W_k(x, x'), \tag{4}$$

where $\lambda_k$ $(k = 1, \cdots, \ell)$ are the kernel hyperparameters and correspond to the variance of the coefficients representing pure epistasic interactions of order $k$, expressed under any orthonormal basis, and $W_k(x, x')$ is the covariance when the prior only consists of pure $k$-th order interactions that are drawn with unit norm and equal probability among any choice of $k$ sites and depends on the Krawtchouk polynomials [2]:

$$W_k(x, x') = \alpha^{-\ell} \sum_{j=0}^{k} (-1)^j (\alpha - 1)^{k-j} \binom{d(x, x')}{j} \binom{\ell - d(x, x')}{k - j}. \tag{5}$$

The $\lambda_k$ are directly related to the variance components in the prior, that is, the proportion of total variance explained by interactions among $k$ site under the prior.

For the Gaussian process regression models presented in this paper, we first infer the hyperparameters $\theta$ (e.g. the $\lambda_k$ for VC regression or the $\delta_p^a$ for the Jenga model) through evidence maximization, that is, we choose $\theta$ that maximize the marginal likelihood (the probability of observing the data given $\theta$). Specifically,

$$\widehat{\theta} = \arg\max_{\theta} \log p(y|\theta) = \arg\max_{\theta} \left[ -\frac{n}{2}\log(2\pi) - \frac{1}{2}\log|K_{BB}(\theta) + E(\theta)| - \frac{1}{2} y^T (K_{BB}(\theta) + E(\theta))^{-1} y) \right], \tag{6}$$

where $n$ is the number of measured sequences. The inferred hyperparameters $\widehat{\theta}$ under different priors provide different biological insights into the data, such as the relative importance of interaction of different orders

(VC regression), and the effect of mutating specific positions or alleles on the predictability of mutational effects (connectedness, Jenga and general product models). Importantly, with the inferred hyperparameters $\widehat{\theta}$, we can then calculate the posterior mean and covariance for any set of sequences using standard results in Eqs 2 and 3.

#### 1.2 Correlation of mutational effects across genetic backgrounds under our priors

In this section, we derive the correlation of mutational effects across backgrounds in the connectedness model and the Jenga model. In fact, this correlation is simply given by their respective kernel functions.

Let $p$ be the position where the focal mutation $a \to a'$ occurs. Assume there are two genotypes $x, x'$ that segregate at positions $D \subset \{1, \cdots, \ell\}$. Let $S$ denote the subset of sites excluding $p$ where the two genotypes are identical. We have $D \cup S \cup \{p\} = \{1, \cdots, \ell\}$. Let $f_{S,D,p}^{x_S,x_D,a}$ denote the fitness of $x$, where $x_S, x_D, a$ symbolize the its genotype at positions $S$, $D$, and $p$ respectively. Then the effect of the focal mutation in the $x$ background can be represented as

$$f_{S,D,p}^{x_S,x_D,a'} - f_{S,D,p}^{x_S,x_D,a}.$$

Similarly, the same mutational effect calculated in the $x'$ background is

$$f_{S,D,p}^{x_S,x'_D,a'} - f_{S,D,p}^{x_S,x'_D,a}.$$

Since both priors assumes uniform zero mean, the covariance between these two quantities is

$$\mathbb{E}\left[(f_{S,D,p}^{x_S,x_D,a'} - f_{S,D,p}^{x_S,x_D,a})(f_{S,D,p}^{x_S,x'_D,a'} - f_{S,D,p}^{x_S,x'_D,a})\right], \tag{7}$$

where the expectation is taken over the distribution over all $f$ under specific priors i.e. connectedness, Jenga or general product priors.

Under the connectedness model or the Jenga model, the covariance in phenotypes is simply given by their respective kernels. Using the fact that the correlation in both models factorizes over positions, this can be re-expressed as

$$\begin{aligned}\mathbb{E}\left[(f_{S,D,p}^{x_S,x_D,a'} - f_{S,D,p}^{x_S,x_D,a})(f_{S,D,p}^{x_S,x'_D,a'} - f_{S,D,p}^{x_S,x'_D,a})\right] =& r_D^{x_D,x'_D} + r_D^{x_D,x'_D} - \left(r_D^{x_D,x'_D} r_p^{a,a'} + r_D^{x_D,x'_D} r_p^{a,a'}\right) \\ =& 2r_D^{x_D,x'_D}(1 - r_p^{a,a'}),\end{aligned} \tag{8}$$

where we have used the generic notation $r_D^{x_D,x'_D}$ to denote the correlation between two genotypes segregating on sites $D$, with genotypes $x_D, x'_D$.

Similarly, we can calculate the variance in the mutational effects $a \to a'$ in the two backgrounds

$$\mathbb{E}\left[(f_{S,D,p}^{x_S,x_D,a'} - f_{S,D,p}^{x_S,x_D,a})^2\right] = \mathbb{E}\left[(f_{S,D,p}^{x_S,x'_D,a'} - f_{S,D,p}^{x_S,x'_D,a})^2\right] = 2(1 - r_p^{a,a'}).$$

Thus, by taking the ratio we have a quantity that measures the correlation in the effect of the mutation $a \to a'$ between the two genotypes $x$ and $x'$

$$\frac{\mathbb{E}\left[(f_{S,D,p}^{x_S,x_D,a'} - f_{S,D,p}^{x_S,x_D,a})(f_{S,D,p}^{x_S,x'_D,a'} - f_{S,D,p}^{x_S,x'_D,a})\right]}{\mathbb{E}\left[(f_{S,D,p}^{x_S,x_D,a'} - f_{S,D,p}^{x_S,x_D,a})^2\right]^{1/2} \mathbb{E}\left[(f_{S,D,p}^{x_S,x'_D,a'} - f_{S,D,p}^{x_S,x'_D,a})^2\right]^{1/2}} = r_D^{x_D,x'_D} \tag{9}$$

In the connectedness model, this becomes

$$\prod_{p \in D}(1 - \delta_p). \tag{10}$$

In the Jenga model, this becomes

$$\prod_{p \in D}(1 - \delta_p^{x_p})(1 - \delta_p^{x'_p}). \tag{11}$$

Last, in the general product kernel, this becomes

$$\prod_{p\in D}(1-\delta_p^{x_p,x'_p}). \tag{12}$$

These correspond closely with Eqs. 3, 5 6. This result is general for any sequence kernel that is derived as a product of site-specific kernels, revealing the implicit assumption that, in this class of models, mutations not only have the same effect on the correlation in the phenotypes across genetic backgrounds, but also have the same effect on the correlation of mutational effects.

#### 1.3 Relationship between the decay factors and the $\gamma$ statistics

In this section, we derive the correlation of mutational effects in different genetic backgrounds at different Hamming distances to draw connections to the $\gamma$ statistics in [4, 5].

##### 1.3.1 Relationship with $\gamma_d$

First, given a fitness landscape $f$, the empirical covariance in the effect of a mutation between genetic backgrounds at Hamming distance $d$ can be written as

$$\frac{1}{C}\sum_p\sum_{a\neq a'}\sum_{|D|=d}\sum_{x'_D\neq x_D}\sum_{x_S}(f_{S,D,p}^{x_S,x_D,a'}-f_{S,D,p}^{x_S,x_D,a})(f_{S,D,p}^{x_S,x'_D,a'}-f_{S,D,p}^{x_S,x'_D,a}), \tag{13}$$

where $C$ is the normalizing constant corresponding to the number of pairs of the same mutations at distance $d$. Using $\widehat{\rho}_d$ to denote the empirical covariance in phenotypes for sequences at Hamming distance $d$, we can expand Eq. 13 and express it as

$$2\widehat{\rho}_d-2\widehat{\rho}_{d+1}.$$

Setting $d=0$, we acquired the empirical variance in the mutational effects

$$2\widehat{\rho}_0-2\widehat{\rho}_1=2-2\widehat{\rho}_1,$$

assuming the variance in phenotypes is normalized such that $\widehat{\rho}_0=1$.

Therefore, the empirical correlation $\gamma_d$ for mutational effect in genetic background with divergence $d$ is

$$\gamma_d=\frac{\widehat{\rho}_d-\widehat{\rho}_{d+1}}{1-\widehat{\rho}_1}.$$

In the special case $d=1$, this becomes

$$\gamma_1=\frac{\widehat{\rho}_2-\widehat{\rho}_1}{1-\widehat{\rho}_1}, \tag{14}$$

which is the $\gamma$ parameter first introduced in [4]. Here we keep the subscript 1 for clarity.

Here, we want to study the expectation of this quantity $\mathbb{E}[\gamma_d(f)]$ when $f$ itself is a random field to examine how different parameterizations of this distribution affect the overall correlation in mutational effects by Hamming distance. Because calculating the expectation of the ratio in Eq. 14 is difficult, here we calculate the expectation of the covariance in the numerator and the variance in the denominator separately and use the ratio $\Gamma_d$ to measure the distance correlation in mutational effects under different priors for $f$.

Taking the expectation of this empirical covariance in Eq. 13 with respect to our prior distribution, we get

$$\begin{aligned}
&\mathbb{E}\left[\frac{1}{C}\sum_p\sum_{a\neq a'}\sum_{|D|=d}\sum_{x'_D\neq x_D}\sum_{x_S}(f_{S,D,p}^{x_S,x_D,a'}-f_{S,D,p}^{x_S,x_D,a})(f_{S,D,p}^{x_S,x'_D,a'}-f_{S,D,p}^{x_S,x'_D,a})\right]\\
=&\frac{1}{C}\sum_p\sum_{a\neq a'}\sum_{|D|=d}\sum_{x'_D\neq x_D}\sum_{x_S}\mathbb{E}\left[(f_{S,D,p}^{x_S,x_D,a'}-f_{S,D,p}^{x_S,x_D,a})(f_{S,D,p}^{x_S,x'_D,a'}-f_{S,D,p}^{x_S,x'_D,a})\right] \qquad (15)\\
=&\frac{1}{C}\sum_p\sum_{a\neq a'}\sum_{|D|=d}\sum_{x'_D\neq x_D}\sum_{x_S}2r_D^{x_D,x'_D}(1-r_p^{a,a'}),
\end{aligned}$$

using Eq. 8 from the previous section. Recall that $r_D^{x_D,x'_D}$ is a generic notation denoting the correlation between two genotypes segregating on sites $D$ under our product kernels. This expectation can be further expressed compactly as

$$2\langle r_D^{x_D,x'_D}\rangle_{|D|=d,x_D\neq x_{D'}} - 2\langle r_{D'}^{x_{D'},x'_{D'}}\rangle_{|D'|=d+1,x_{D'}\neq x'_{D'}}, \tag{16}$$

where the first term denotes the average taken over all subsets $D$ of size $d$ and all pairs of sequences of length $d$, such that $x_D \neq x'_D$. The second term is similar, except that $D'$ contains $d+1$ positions.

Setting $d = 0$, we can get the the expected empirical variance of all mutational effects

$$2 - 2\langle r_p^{a,a'}\rangle_{p,a\neq a'}. \tag{17}$$

We can take the ratio of Eq. 16 and Eq. 17 and arrive at $\Gamma_d$, which characterizes the similarity in mutational effects separated by Hamming distance $d$ for any product kernel prior.

$$\Gamma_d = \frac{\langle r_D^{x_D,x'_D}\rangle_{|D|=d,x_D\neq x_{D'}} - \langle r_{D'}^{x_{D'},x'_{D'}}\rangle_{|D'|=d+1,x_{D'}\neq x'_{D'}}}{1 - \langle r_p^{a,a'}\rangle_{p,a\neq a'}}. \tag{18}$$

$\Gamma_d$ is similar to the $\gamma_d$ statistic introduced earlier. The key difference is that $\gamma_d$ is a statistic of a realized fitness landscape $f$, whereas $\Gamma_d$ represents the expectation of numerator and denominator in Eq. 18 of their counterparts in Eq. 14 under the prior distribution. To parallel the expression in Eq. 14, we can express $\Gamma_1$ as

$$\Gamma_1 = \frac{\rho_1 - \rho_2}{1 - \rho_1}. \tag{19}$$

And more generally,

$$\Gamma_d = \frac{\rho_d - \rho_{d+1}}{1 - \rho_1}. \tag{20}$$

Note that $\rho_d = \langle r_D^{x_D,x'_D}\rangle_{|D|=d,x_D\neq x_{D'}}$ is an average over expectations. Using the definition of our kernels, we can derive explicit expressions of Eq. 20.

First, in the connectedness model, we have

$$\rho_d = \langle r_D^{x_D,x'_D}\rangle_{|D|=d,x_D\neq x_{D'}} = \frac{1}{\binom{\ell}{d}} e_d\left(1-\delta_1, \cdots, 1-\delta_\ell\right), \tag{21}$$

where $e_d()$ denotes the elementary symmetric polynomial of degree $d$.

$$e_d(x_1, x_2, \ldots, x_n) = \sum_{1\leq b_1<b_2<\cdots<b_d\leq n} x_{b_1}x_{b_2}\cdots x_{b_1}.$$

Consequently, we find

$$\rho_1 = \langle 1 - \delta_p\rangle_p.$$

In the Jenga model, we have

$$\rho_d = \frac{1}{\binom{\ell}{d}}\langle e_d\left((1-\delta_1^{a_1})(1-\delta_1^{a'_1}), \cdots, (1-\delta_\ell^{a_\ell})(1-\delta_\ell^{a'_\ell})\right)\rangle_{a_1\neq a'_1,\cdots,a_\ell\neq a'_\ell} \tag{22}$$

and

$$\rho_1 = \langle(1-\delta_p^a)(1-\delta_p^{a'})\rangle_{p,a\neq a'}.$$

Finally, for the general product kernel

$$\rho_d = \frac{1}{\binom{\ell}{d}}\langle e_d\left((1-\delta_1^{a_1,a'_1}), \cdots, (1-\delta_\ell^{a_\ell,a'_\ell})\right)\rangle_{a_1\neq a'_1,\cdots,a_\ell\neq a'_\ell}, \tag{23}$$

while

$$\rho_1 = \langle(1-\delta^{a_p,a'_p})\rangle_{p,a\neq a'}.$$

Using these expressions for $\rho_2$ and $\rho_1$, we can see that $\Gamma_1 = \frac{\rho_1 - \rho_2}{1-\rho_1}$ does not have a simple relationship with the decay factors under different priors.

However, we can show that despite this, $\Gamma_1$ does decrease monotonically with any $\delta$, such that increasing any decay factor results in the decrease in predictability of all other mutations. We show this for the most general product kernel, by first expanding both $\rho_1$ and $\rho_2$

$$\rho_1 = \frac{1}{\ell\binom{\alpha}{2}} \sum_p \sum_{a_p < a'_p} (1 - \delta_p^{a_p, a'_p}) = 1 - \frac{1}{\ell\binom{\alpha}{2}} \sum_p \sum_{a_p < a'_p} \delta_p^{a_p, a'_p},$$

$$\begin{aligned}
\rho_2 =& \frac{1}{\binom{\ell}{2}\binom{\alpha}{2}^2} \sum_{p<q} \sum_{a_p < a'_p} \sum_{a_q < a'_q} \left(1 - \delta_p^{a_p, a'_p}\right)\left(1 - \delta_q^{a_q, a'_q}\right) \\
=& \frac{1}{\binom{\ell}{2}\binom{\alpha}{2}^2} \left[ \binom{\ell}{2}\binom{\alpha}{2}^2 - (\ell - 1)\binom{\alpha}{2} \sum_p \sum_{a_p < a'_p} \delta_p^{a_p, a'_p} + \sum_{p<q} \sum_{\substack{a_p < a'_p \\ a_q < a'_q}} \delta_p^{a_p, a'_p} \delta_q^{a_q, a'_q} \right] \\
=& 1 - \frac{2}{\ell\binom{\alpha}{2}} \sum_p \sum_{a_p < a'_p} \delta_p^{a_p, a'_p} + \frac{1}{\binom{\ell}{2}\binom{\alpha}{2}^2} \sum_{p<q} \sum_{\substack{a_p < a'_p \\ a_q < a'_q}} \delta_p^{a_p, a'_p} \delta_q^{a_q, a'_q},
\end{aligned}$$

and expressing $\Gamma_1$ as

$$\begin{aligned}
\Gamma_1 =& \frac{\rho_1 - \rho_2}{1 - \rho_1} \\
=& \frac{\left(1 - \frac{1}{\ell\binom{\alpha}{2}} \sum_p \sum_{a_p < a'_p} \delta_p^{a_p, a'_p}\right) - \left(1 - \frac{2}{\ell\binom{\alpha}{2}} \sum_p \sum_{a_p < a'_p} \delta_p^{a_p, a'_p} + \frac{1}{\binom{\ell}{2}\binom{\alpha}{2}^2} \sum_{p<q} \sum_{\substack{a_p < a'_p \\ a_q < a'_q}} \delta_p^{a_p, a'_p} \delta_q^{a_q, a'_q}\right)}{\frac{1}{\ell\binom{\alpha}{2}} \sum_p \sum_{a_p < a'_p} \delta_p^{a_p, a'_p}} \\
=& \frac{\frac{1}{\ell\binom{\alpha}{2}} \sum_p \sum_{a_p < a'_p} \delta_p^{a_p, a'_p} - \frac{1}{\binom{\ell}{2}\binom{\alpha}{2}^2} \sum_{p<q} \sum_{\substack{a_p < a'_p \\ a_q < a'_q}} \delta_p^{a_p, a'_p} \delta_q^{a_q, a'_q}}{\frac{1}{\ell\binom{\alpha}{2}} \sum_p \sum_{a_p < a'_p} \delta_p^{a_p, a'_p}} \\
=& \frac{\sum_p \sum_{a_p < a'_p} \delta_p^{a_p, a'_p} - \frac{2}{(\ell - 1)\binom{\alpha}{2}} \sum_{p<q} \sum_{\substack{a_p < a'_p \\ a_q < a'_q}} \delta_p^{a_p, a'_p} \delta_q^{a_q, a'_q}}{\sum_p \sum_{a_p < a'_p} \delta_p^{a_p, a'_p}}.
\end{aligned} \tag{24}$$

Now, can take the derivative with respect to $\delta_p^{a_p, a'_p}$:

$$
\begin{aligned}
\frac{\partial \Gamma_1}{\partial \delta_p^{a_p,a_p'}} =& \frac{\left(1-\frac{2}{(\ell-1)\binom{\alpha}{2}}\sum_{q\neq p}\sum_{a_q<a_q'}\delta_q^{a_q,a_q'}\right)\left(\sum_m\sum_{a_m<a_m'}\delta_m^{a_m,a_m'}\right)}{\left(\sum_m\sum_{a_m<a_m'}\delta_m^{a_m,a_m'}\right)^2} \\
&-\frac{\left(\sum_m\sum_{a_m<a_m'}\delta_m^{a_m,a_m'}-\frac{2}{(\ell-1)\binom{\alpha}{2}}\sum_{q<m}\sum_{\substack{a_m<a_m'\\a_q<a_q'}}\delta_m^{a_m,a_m'}\delta_q^{a_q,a_q'}\right)}{\left(\sum_m\sum_{a_m<a_m'}\delta_m^{a_m,a_m'}\right)^2} \\
=& \frac{-\left(\sum_m\sum_{a_m<a_m'}\delta_m^{a_m,a_m'}\right)\left(\frac{2}{(\ell-1)\binom{\alpha}{2}}\sum_{q\neq p}\sum_{a_q<a_q'}\delta_q^{a_q,a_q'}\right)+\frac{2}{(\ell-1)\binom{\alpha}{2}}\sum_{q<m}\sum_{\substack{a_m<a_m'\\a_q<a_q'}}\delta_m^{a_m,a_m'}\delta_q^{a_q,a_q'}}{\left(\sum_m\sum_{a_m<a_m'}\delta_m^{a_m,a_m'}\right)^2} \\
=& -\frac{2}{(\ell-1)\binom{\alpha}{2}}\frac{\sum_{q\neq p,m}\sum_{\substack{a_m<a_m'\\a_q<a_q'}}\delta_m^{a_m,a_m'}\delta_q^{a_q,a_q'}-\sum_{q<m}\sum_{\substack{a_m<a_m'\\a_q<a_q'}}\delta_m^{a_m,a_m'}\delta_q^{a_q,a_q'}}{\left(\sum_m\sum_{a_m<a_m'}\delta_m^{a_m,a_m'}\right)^2},
\end{aligned}
$$

where

$$
\sum_{q\neq p,m}\sum_{\substack{a_m<a_m'\\a_q<a_q'}}\delta_m^{a_m,a_m'}\delta_q^{a_q,a_q'}=\sum_{q\neq p}\sum_{a_q<a_q'}(\delta_q^{a_q,a_q'})^2+\sum_{\substack{q<m\\q\neq p\\m\neq p}}\sum_{\substack{a_m<a_m'\\a_q<a_q'}}\delta_m^{a_m,a_m'}\delta_q^{a_q,a_q'}+\sum_{q<m}\sum_{\substack{a_m<a_m'\\a_q<a_q'}}\delta_m^{a_m,a_m'}\delta_q^{a_q,a_q'},
$$

so

$$
\frac{\partial \Gamma_1}{\partial \delta_p^{a_p,a_p'}}=-\frac{2}{(\ell-1)\binom{\alpha}{2}}\frac{\sum_{q\neq p}\sum_{a_q<a_q'}(\delta_q^{a_q,a_q'})^2+\sum_{\substack{q<m\\q\neq p\\m\neq p}}\sum_{\substack{a_m<a_m'\\a_q<a_q'}}\delta_m^{a_m,a_m'}\delta_q^{a_q,a_q'}}{\left(\sum_m\sum_{a_m<a_m'}\delta_m^{a_m,a_m'}\right)^2}. \tag{25}
$$

Since $\delta_p^{a_p,a_p'}\geq 0$, $\frac{\partial\Gamma_1}{\partial\delta_p^{a_p,a_p'}}=0$ if and only if $\delta_q^{a_q,a_q'}=0$ for all $q\neq p, a_q, a_q'$ combinations. Moreover, for any set of valid $\delta_p^{a_p,a_p'}>0$, $\frac{\partial\Gamma_1}{\partial\delta_p^{a_p,a_p'}}<0$ and thus $\Gamma_1$ monotonically decreases with $\delta_p^{a_p,a_p'}$.

###### 1.3.2 Generalization of $\gamma$ and its relationship to the decay factors

Despite this complex but monotonic relationship between $\Gamma_1$ and $\delta$, there is indeed a generalization of the $\gamma$ statistics that is directly related to our decay factors. For a position $q$ and a mutation $b\to b'$, the empirical covariance in mutational effects between genotypes segregated by the mutation $b\to b'$ or $b'\to b$ at position $q$ can be written as

$$
\frac{1}{C}\sum_{p\neq q}\sum_{a\neq a'}\sum_{S\not\ni q,p}\sum_{x_S}\left[(f_{S,q,p}^{x_S,b,a'}-f_{S,q,p}^{x_S,b,a})(f_{S,q,p}^{x_S,b',a'}-f_{S,q,p}^{x_S,b',a})\right]. \tag{26}
$$

Let $\gamma_q^{b,b'}$ be the empirical correlation in the effects of mutations across genetic backgrounds differing at position $q$ with alleles $b$ and $b'$, which can be obtained by dividing the empirical covariance by the variance in a given fitness landscape $f$:

$$
\gamma_q^{b,b'}=\frac{\frac{1}{C}\sum_{p\neq q}\sum_{a\neq a'}\sum_{S\not\ni q,p}\sum_{x_S}\left[(f_{S,q,p}^{x_S,b,a'}-f_{S,q,p}^{x_S,b,a})(f_{S,q,p}^{x_S,b',a'}-f_{S,q,p}^{x_S,b',a})\right]}{\frac{1}{C'}\sum_{p\neq q}\sum_{a\neq a'}\sum_{x_S}\sum_b(f_{S,q,p}^{x_S,b,a'}-f_{S,q,p}^{x_S,b,a})^2}, \tag{27}
$$

where $C$ and $C'$ are the appropriate normalizing constants.

Taking expectations of the numerator and the denominator with respect to our priors, we get a new quantity, namely $\Gamma_q^{b,b'}$, which provides a measurement of the average correlation in mutational effects across backgrounds separated by the $b\to b$ mutation. Specifically, we find

$$\Gamma_q^{b,b'} = \frac{\langle 2r_q^{b,b'} - 2r_q^{b,b'} r_p^{a,a'} \rangle_{p\neq q, a\neq a'}}{\langle 2 - 2r_p^{a,a'} \rangle_{p\neq q, a\neq a'}} = \frac{r_q^{b,b'}(1 - \langle r_p^{a,a'} \rangle_{p\neq q, a\neq a'})}{1 - \langle r_p^{a,a'} \rangle_{p\neq q, a\neq a'}} = r_q^{b,b'}, \tag{28}$$

which is equal to $1 - \delta_q$ in the connectedness model, $(1 - \delta_q^b)(1 - \delta_q^{b'})$ in the Jenga model, and $1 - \delta_q^{b,b'}$ in the general product model.

Therefore, we can see that the decay factors in our kernels are more directly related to the empirical measurement of epistatic effect of a particular mutation at a locus on other loci given by $\gamma_q^{b,b'}$ statistic, instead of the more coarse-grained correlation in terms of Hamming distance $\gamma_d$. Note that Eq. 27 is identical to Eq. S1-9 in in Bank et al. [5] as an intermediate result for deriving $\gamma_d$. Last, Eq. 27 and Eq. S1-9 in in Bank et al. are difficult to evaluate in practice when the genotype space is large as one needs to enumerate through mutations. Here, we present a more concise formula expressed in terms of the correlation in phenotypes to allow practical computation in large datasets

$$\gamma_q^{b,b'} = \frac{\widehat{\rho}_q^{b,b'} - \langle \widehat{\rho}_{pq}^{aa',bb'} \rangle_{p\neq q, a\neq a'}.}{1 - \langle \widehat{\rho}_p^{a,a'} \rangle_{p\neq q, a\neq a'}}. \tag{29}$$

Recall that $\widehat{\rho}_q^{b,b'}$ denotes the empirical correlation between genotypes having alleles $b$ and $b'$ at position $q$, and $\widehat{\rho}_{pq}^{aa',bb'}$ is the version when they segregating at loci $p$ and $q$, with genotypes $aa'$ and $bb'$. This can be derived by simply expanding the numerator and denominator of Eq. 27.

#### 1.4 Relationship between the connectedness model $\delta_p$ and $\mu_p$

In this section, we will establish the relationship between the connectedness prior used in the connectedness regression and the original connectedness model proposed by Reddy and Desai [6]. Specifically, we will show the relationship between our parameterization in terms of the decay factors $\delta_p$ and the original parameter $\mu_p$, revealed in Eq. 4 in the main text.

We first derive the kernel function for the prior distribution under the original connectedness model parametrized by $\mu_p$. The original connectedness model assigns a parameter $\mu_p$ to each position $i$ and assumes that the magnitude of the interaction terms involving any subset of loci is proportional to the product of their individual parameters. That is, for a three-way interaction involving loci $i, j, k$, the interaction term $f_{ijk}^2 \propto \mu_p \mu_j \mu_k$.

We further assume that all interaction terms are Gaussian with zero mean (including the constant term) and that these coefficients are expressed with respect to an orthonormal basis (e.g. the Walsh-Fourier basis) for the space of all sequence-function relationships. First, consider the Walsh-Fourier basis for a single position. The projection matrix $P_{\text{con}}$ onto the space of constant functions is

$$P_{\text{con}}(a, a') = \frac{1}{\alpha}. \tag{30}$$

The projection matrix $P_{\text{con}}$ onto the residual subspace of additive functions $P_{\text{add}}$ is $I - P_{\text{con}}$. Therefore,

$$P_{\text{add}}(a, a') = \begin{cases} \frac{\alpha-1}{\alpha} & a = a' \\ -\frac{1}{\alpha} & a \neq a. \end{cases} \tag{31}$$

Then the projection matrix onto the subspace spanned by interactions among a subset of positions $U$ can be derived by taking tensor products of Eq. 30 and Eq. 31.

$$P_U(x, x') = \alpha^{-\ell} \prod_{p\in U: x_p = x'_p} (\alpha - 1) \prod_{p\in U: x_p \neq x'_p} (-1). \tag{32}$$

Under the assumption that the interaction coefficients are i.i.d with zero mean and $f_{ijk}^2 \propto \mu_p \mu_j \mu_k$, the covariance when only considering all possible interactions among positions in $U$ is

$$\mathbb{E}[f(x)f(x')] = \prod_{p\in U} \mu_p \times P_U(x, x').$$

Considering the interactions among all subsets of sites, the covariance becomes

$$\begin{aligned}\mathbb{E}[f(x)f(x')] &= \alpha^{-\ell}\sum_{U}\prod_{p\in U}\mu_p \prod_{p\in U: x_p = x'_p}(\alpha-1)\prod_{p\in U: x_p\neq x'_p}(-1)\\ &= \alpha^{-\ell}\prod_{p:x_p=x'_p}(1+\mu_p(\alpha-1))\prod_{p:x_p\neq x'_p}(1-\mu_p).\end{aligned}$$

Normalizing by the variance and re-scaling by $\sigma^2$, we get the correlation matrix

$$k(x,x') = \sigma^2 \prod_{p\in U: x_p\neq x'_p}\frac{1-\mu_p}{1+\mu_p(\alpha-1)}. \tag{33}$$

Comparing this with the expression of the connectedness kernel

$$k(x,x') = \sigma^2\prod_{p:x_p\neq x'_p}(1-\delta_p),$$

we arrive at the results in Eq. 4 in the main text:

$$\delta_p = 1-\frac{1-\mu_p}{1+\mu_p(\alpha-1)} = \frac{1+\mu_p(\alpha-1)-(1-\mu_p)}{1+\mu_p(\alpha-1)} = \frac{\mu_p\alpha}{1+\mu_p(\alpha-1)}. \tag{34}$$

#### 1.5 Relationship between the Jenga $\delta_p^a$ and the automatic relevance determination kernel length scales

In the main text, we mentioned the connection with the Jenga kernel in Eq. 5 and the classical automatic relevance determination (ARD) kernel over a space with $n$ different dimensions

$$k(x,x') = \sigma^2 \exp\left(-\frac{1}{2}\sum_{i=1}^{n}\frac{(x_i-x'_i)^2}{l_i^2}\right). \tag{35}$$

The relation between these two kernels can be seen by first converting the input genotypes using one-hot encoding, such that each genotype is transformed to an $n=\alpha\ell$ dimensional vector, with each entry equal to the evaluation of an indicator function for allele $a$ at a position $p$, i.e.

$$x = [\mathbf{1}(x_p=a)]_{p=1,\cdots,\ell;a=1,\cdots,\alpha}.$$

Now assigning a length scale parameter $l_{p,a}$ to allele $a$ at position $p$ in the continuous $\mathbb{R}^{\alpha\times\ell}$ space, we can apply Eq. 35 to two genotypes $x$ and $x'$:

$$k(x,x') = \sigma^2\prod_{p}\prod_{a}\exp\left(-\frac{1}{2}\frac{\left(\mathbf{1}(x_p=a)-\mathbf{1}(x'_p=a)\right)^2}{l_{p,a}^2}\right).$$

Thus, for each position $p$, the kernel evaluates to

$$\prod_{a}\exp\left(-\frac{1}{2}\frac{\left(\mathbf{1}(x_p=a)-\mathbf{1}(x'_p=a)\right)^2}{l_{p,a}^2}\right) = \begin{cases}1 & \text{if } x_p = x'_p\\ \exp\left(-\frac{1}{2l_{p,a}^2}\right)\times\exp\left(-\frac{1}{2l_{p,a'}^2}\right) & \text{if } x_p\neq x'_p,\end{cases}$$

where $a$ and $a'$ are the alleles for $x$ and $x'$ at position $p$, respectively. Comparing with the Jenga kernel

$$k(x,x') = \sigma^2\prod_{p:x_p\neq x'_p}(1-\delta_p^a)(1-\delta_p^{a'}).$$

Thus, we have this relationship

$$l_{p,a}^2 = -\frac{1}{2\log(1-\delta_p^a)}. \tag{36}$$

#### 1.6 Generalization of the Jenga and connectedness models for $\delta > 1$

In the main text, we described the connectedness and Jenga kernels for $0 < \delta_p < 1$ and $0 < \delta_p^a < 1$, in which the resulting kernel is equivalent to some forms of the Automatic Relevance Determination (ARD) kernel on the one-hot encoded sequences. In this section, we generalize it for $\delta_p > 1$, enabling negative correlations between pairs of sequences. This is possible because kernel only needs to be positive definite when defined in the complete and discrete sequence space as opposed to $\mathbb{R}^N$. However, there are two constraints on the values $\delta_p^a$ can take. First, all $\delta_p^a$'s must be either smaller or larger than 1 for every allele $a$ within a single site $p$. Second, not all $\delta_p^a > 1$ result in valid positive definite kernel matrices. Under these two conditions, the Jenga kernel can be generalized as follows:

$$k(x, x') = \prod_{p: x_p \neq x'_p} \mathrm{sign}\left(1 - \delta_p^a\right) |1 - \delta_p^a||1 - \delta_p^{a'}|. \tag{37}$$

To derive the bounds on $\delta_p^a$, we resort to a different formulation of the Jenga kernel. As it is a product kernel, it is sufficient to show its properties for a single-site kernel ($\ell = 1$) with $\alpha$ different alleles. We first define a probability distribution $\pi_a$ over all $\alpha$ possible alleles, such that $\sum_{a=1}^{\alpha} \pi_a = 1$. Let $D_\pi$ denote the diagonal matrix such that $(D_\pi)_{a,a} = \pi_a$. We can then decompose any $\alpha$-dimensional function $f$ in two $D_\pi$-orthogonal components, corresponding to the constant subspace and its complement ($f = f_0 + f_1$), which are defined by their $D_\pi$-orthogonal projection matrices $P_{\text{con}}^{(\pi)} = 11^T D_\pi$ and $P_{\text{add}}^{(\pi)} = I - 11^T D_\pi$, respectively.

We define our prior distribution by sampling a vector $b$ from a zero-mean Gaussian distribution with covariance matrix $D_\pi^{-1}$ and projecting it onto them onto the different subspaces ($f_{\text{con}} = P_{\text{con}}^{(\pi)} b$ and $f_{\text{add}} = P_{\text{add}}^{(\pi)} b$). As both components are obtained by a linear combination of a Gaussian distribution, they also a follow a Gaussian distribution defined by their covariance matrices $f_{\text{con}} \sim N(0, K_{\text{con}}^{(\pi)})$ and $f_{\text{add}} = N(0, K_{\text{add}}^{(\pi)})$, respectively:

$$K_{\text{con}}^{(\pi)} = P_{\text{con}}^{(\pi)} D_\pi^{-1} (P_{\text{con}}^{(\pi)})^T = 11^T D_\pi D_\pi^{-1} (11^T D_\pi)^T = 11^T D_\pi 11^T = 11^T.$$

$$\begin{aligned} K_{\text{add}}^{(\pi)} &= P_{\text{add}}^{(\pi)} D_\pi^{-1} (P_{\text{add}}^{(\pi)})^T = (I - 11^T D_\pi) D_\pi^{-1} (I - 11^T D_\pi)^T = (D_\pi^{-1} - 11^T)(I - 11^T D_\pi)^T = \\ &= D_\pi^{-1} - D_\pi^{-1} D_\pi 11^T - 11^T + 11^T D_\pi 11^T = D_\pi^{-1} - 11^T - 11^T + 11^T = D_\pi^{-1} - 11^T. \end{aligned}$$

We derive the covariance matrix for $f \sim N(0, K)$ as the weighted sum of the covariances of the two components with relative weights 1 and $\mu$ as $K = K_{\text{con}}^{(\pi)} + \mu K_{\text{add}}^{(\pi)} = 11^T + \mu(D_\pi^{-1} - 11^T)$, with entry-wise formula given by:

$$k(x, x') = \begin{cases} 1 + \mu \frac{1 - \pi_x}{\pi_x} & \text{if } x = x' \\ 1 - \mu & \text{otherwise.} \end{cases}$$

Lets now consider the correlation matrix instead:

$$k(x, x') = \begin{cases} 1 & \text{if } x = x' \\ \frac{1 - \mu}{\sqrt{1 + \mu \frac{1 - \pi_x}{\pi_x}} \sqrt{1 + \mu \frac{1 - \pi_{x'}}{\pi_{x'}}}} & \text{otherwise.} \end{cases}$$

Under a uniform probability distribution over the alleles $\pi = \frac{1}{\alpha}$, the off-diagonal terms simplify to the connectedness model, given by $\frac{1-\mu}{1+(\alpha-1)\mu} = 1 - \delta_p$. The maximal decay rate $\delta_p$ is achieved as $\lim_{\mu \to \infty} \frac{1-\mu}{1+(\alpha-1)\mu} = -\frac{1}{\alpha - 1}$ and thus:

$$\delta_p \leq \frac{\alpha}{\alpha - 1}. \tag{38}$$

Note that the sign of the off-diagonal terms depends only on $\mu$, which can be site-specific, but not on $\pi_a$. Therefore all $k(x, x')$ for $x \neq x'$ must be either positive or negative at the same time. Note that this kernel can also be written as a product of allele-specific decay factors, showing that they correspond to the same family of kernels.

$$\frac{1-\mu}{\sqrt{1+\mu\frac{1-\pi_a}{\pi_a}}\sqrt{1+\mu\frac{1-\pi_{a'}}{\pi_{a'}}}} = \text{sign}(1-\mu)\sqrt{\frac{|1-\mu|}{1+\mu\frac{1-\pi_a}{\pi_a}}}\sqrt{\frac{|1-\mu|}{1+\mu\frac{1-\pi_{a'}}{\pi_{a'}}}} = \text{sign}(1-\mu)|1-\delta^a||1-\delta^{a'}|.$$

Importantly, by construction, as long as $D_\pi^{-1}$ is positive definite, which is true for any valid $\pi$, so are the covariance and correlation matrices, providing a way to define the bounds on valid $\delta^a$'s. Lets start by defining $\pi_a$ as a function of $\delta^a$:

$$\begin{aligned}
\frac{|1-\mu|}{1-\mu+\frac{\mu}{\pi_a}} &= (1-\delta^a)^2 \\
|1-\mu| &= (1-\mu)(1-\delta^a)^2 + \frac{\mu}{\pi_a}(1-\delta^a)^2 \\
|1-\mu| - (1-\mu)(1-\delta^a)^2 &= \frac{\mu}{\pi_a}(1-\delta^a)^2 \\
\pi_a = \frac{\mu(1-\delta^a)^2}{|1-\mu|-(1-\mu)(1-\delta^a)^2} &= \frac{\mu}{1-\mu}\frac{(1-\delta^a)^2}{\frac{|1-\mu|}{1-\mu}-(1-\delta^a)^2} = \frac{\mu}{1-\mu}\frac{(1-\delta^a)^2}{\text{sign}(1-\delta^a)-(1-\delta^a)^2}.
\end{aligned}$$

We know $\sum_a^\alpha \pi_a = 1$

$$\sum_a^\alpha \pi_a = \sum_a^\alpha \frac{\mu}{1-\mu}\frac{(1-\delta^a)^2}{\text{sign}(1-\delta^a)-(1-\delta^a)^2} = \frac{\mu}{1-\mu}\sum_a^\alpha \frac{(1-\delta^a)^2}{\text{sign}(1-\delta^a)-(1-\delta^a)^2} = 1,$$

which allow us to derive $\mu$ as a function of $\delta^a$'s:

$$\mu = \frac{1}{1+\sum_{a=1}^\alpha \frac{(1-\delta^a)^2}{\text{sign}(1-\delta^a)-(1-\delta^a)^2}}.$$

Thus, any set of $\delta^a$'s that result in $\mu \geq 0$ provides a positive-semidefinite kernel, which is true whenever

$$1+\sum_a^\alpha \frac{(1-\delta^a)^2}{\text{sign}(1-\delta^a)-(1-\delta^a)^2} \geq 0.$$

Note that this is always true for $0<\delta^a<1$, as all $\frac{(1-\delta^a)^2}{1-(1-\delta^a)^2} > 0$ but $\delta^a > 1$ only for a subset of $\delta^a$ given by:

$$\begin{aligned}
1+\sum_a^\alpha \frac{(1-\delta^a)^2}{-1-(1-\delta^a)^2} &\geq 0 \\
1-\sum_a^\alpha \frac{(1-\delta^a)^2}{1+(1-\delta^a)^2} &\geq 0 \\
\sum_{a=1}^\alpha \frac{(1-\delta^a)^2}{1+(1-\delta^a)^2} &\leq 1
\end{aligned} \tag{39}$$

#### 1.7 Variance component computation

We can decompose the space of fitness function vectors $\mathbb{R}^{\alpha^\ell}$ into orthogonal subspaces $V_U$ that represent interactions among the positions in $U$. For a vector $v \in V_U$, the values $v_x$ are the same for all $x$ that have the same subsequence on $U$. The projection matrix into $V_U$ is as follows

$$P_U(x,x') = \alpha^{-\ell}\prod_{\substack{p\in U\\ x_p=x'_p}}(\alpha-1)\prod_{\substack{p\in U\\ x_p\neq x'_p}}(-1)$$

See [2, 7] for more details. Let $V_k$ be the direct sum of $V_U$ for all size $k$ sets of positions $U$. The subspace $V_k$ represents $k^{th}$ order interactions. The orthogonality of the subspaces $V_U$ imply that the projection matrix into $V_k$ is the sum $P_k = \sum_{U:|U|=k} P_U$.

We can understand the geometry of a fitness function vector $f$ by writing it as a sum of vectors in these subspaces:

$$f = \sum_U P_U f \quad \text{or} \quad f = \sum_{k=0}^{\ell} P_k f.$$

The orthogonality of the subspaces implies that

$$f^T f = \sum_{U \in \mathcal{P}([\ell])} (P_U f)^T P_U f = \sum_{k=0}^{\ell} (P_k f)^T P_k f.$$

Such decompositions help us understand how much of the variance of a function drawn from a distribution is due to various subsets of positions or orders of interaction. Let

$$\nu_U = \mathbb{E}[(P_U f)^T (P_U f)] \quad \text{and} \quad \nu_k = \mathbb{E}[(P_k f)^T (P_k f)]$$

be the expected variance due to positions in $U$ or due to $k^{th}$ order interactions. We define the $k^{th}$ *order variance component* as the ratio of the expected variance due to $k^{th}$ order interactions and the total variance,

$$\bar{\nu_k} = \frac{\nu_k}{\mathbb{E}[f^T f]}.$$

We compute the variance components for arbitrary kernels and homoskedastic product kernels.

**Theorem 1.** *Let $f \sim N(0, K)$. Then*

$$\bar{\nu}_k = \frac{\sum_{x,x'} K_{x,x'} \mathcal{K}_k(d(x, x'); \ell, \alpha)}{\text{tr}(K)},$$

*where $d(x, x')$ denotes the Hamming distance between $x$ and $x'$ and $\mathcal{K}_k(d; \ell, \alpha) = \sum_{j=0}^{k} (-1)^j (\alpha-1)^{k-j} \binom{d}{j}\binom{\ell-d}{k-j}$ is the Krawtchouk polynomial.*

In the following theorem $e_k$ denotes the elementary symmetric polynomial of order $k$, the sum of all $k$-way products.

**Theorem 2.** *Let $f \sim N(0, K)$ where $K = \bigotimes_{p=1}^{\ell} K^p$ is a product kernel with each $K^p$ positive definite and ones on the diagonal, $K^p_{x,x} = 1$ for all sequences $x$. Then*

$$\bar{\nu}_k = \left(\prod_{p=1}^{\ell} \zeta_p\right) e_k \left(\frac{1}{\zeta_1} - 1, \ldots, \frac{1}{\zeta_\ell} - 1\right)$$

*where $\zeta_p$ is the average value in $K^p$, $\zeta_p = \frac{1}{\alpha^2} \sum_{c,c'} K^p_{c,c'}$.*

**Corollary 1.** *Let $f \sim N(0, K)$ where $K$ is a connectedness kernel with parameters $\delta_p$. Then*

$$\bar{\nu}_k = \alpha^{-\ell} \left(\prod_{p=1}^{\ell} (\alpha - (\alpha-1)\delta_p)\right) e_k \left(\frac{(\alpha-1)\delta_1}{\alpha - (\alpha-1)\delta_1}, \ldots, \frac{(\alpha-1)\delta_\ell}{\alpha - (\alpha-1)\delta_\ell}\right)$$

**Theorem 3.** *Let $\bar{\nu} = (\bar{\nu}_0, \bar{\nu}_1, \ldots \bar{\nu}_\ell)$ be a vector of non-negative values that sums to one. Then $\bar{\nu}$ are the variance components for some connectedness model if and only if the polynomial*

$$\sum_{k=0}^{\ell} (-1)^k \bar{\nu}_k z^{\ell-k}$$

*has $\ell$ positive roots.*

The same statement applies for general product kernels. Since the connectedness kernels are a subclass of general product kernels, one direction of the if and only if is trivial. To see the other direction (i.e. if $\bar{v}$ are the variance components for some general product kernel, then the polynomial has positive roots), we can follow the proof replacing $\frac{(\alpha-1)\delta_p}{\alpha-(\alpha-1)\delta_p}$ with $\frac{1}{\zeta_p}-1$ throughout. Since each $K^p$ is positive definite, $\zeta_p < 1$ meaning the roots $\frac{1}{\zeta_p}-1$ are positive. This logic also illustrates that there may be infinitely many general product kernels that achieve a particular vector of expected variance components; the variance components are determined by the values $\zeta_p$ and there are usually many ways to construct a positive definite matrix with ones on the diagonal with a particular mean $\zeta_p$. (One exception is the kernel with mean $1/\alpha$, which can only be achieved when $K^p$ is the identity.)

##### 1.7.1 Proof of theorems

**Lemma 1.** *Let $f \sim N(0, K)$.*

$$\nu_U = \sum_{x,x'} (P_U)_{x,x'} K_{x,x'}.$$

*Proof.* Since $P_U$ is a symmetric projection matrix $P_U^T P_U = P_U$. Observe that

$$\nu_U = \mathbb{E}[(P_U f)^T (P_U f)] = \mathbb{E}\left[(f^T P_U^T P_U f)\right] = \mathbb{E}\left[\sum_{x,x'} f_x f_{x'} (P_U^T P_U)_{x,x'}\right] = \sum_{x,x'} (P_U)_{x,x'} K_{x,x'}.$$

□

*Proof.* (of Theorem 1). The orthogonality of the subspaces $V_U$ implies that $\nu_k = \sum_{U:|U|=k} \nu_U$ and $\mathbb{E}[f^T f] = \sum_{k=0}^{\ell} \nu_k$. We first compute $\nu_k$. Let $d_U(x,x')$ denote the Hamming distance of $x[U]$ and $x'[U]$, $|\{p \in U : x_p \neq x'_p\}|$ Lemma 1 implies

$$\nu_U = \alpha^{-\ell} \sum_{x,x'} K_{x,x'} \left( \prod_{\substack{p\in U \\ x_p = x'_p}} (\alpha - 1) \prod_{\substack{p \in U \\ x_p \neq x'_p}} (-1) \right) = \alpha^{-\ell} \sum_{x,x'} K_{x,x'} \left( (\alpha-1)^{|U|-d_U(x,x')} (-1)^{d_U(x,x')} \right). \tag{40}$$

Since $\nu_k = \sum_{|U|=k} \nu_k$,

$$\nu_k = \alpha^{-\ell} \sum_{x,x'} K_{x,x'} \left( \sum_{U:|U|=k} \left( (\alpha-1)^{|U|-d_U(x,x')} (-1)^{d_U(x,x')} \right) \right)$$

Note that the value in the parentheses depends only on the Hamming distance between $x$ and $x'$. Given that $x, x'$ are Hamming distance $d$, there are $\binom{d}{j}\binom{\ell-d}{k-j}$ sets $U$ of size $k$ where $x$ and $x'$ disagree at $j$ positions in $U$, i.e. $d_U(x,x') = j$. This yields

$$\sum_{U:|U|=k} \left( (\alpha-1)^{|U|-d_U(x,x')} (-1)^{d_U(x,x')} \right) = \sum_{j=0}^{k} (-1)^j (\alpha-1)^{k-j} \binom{d(x,x')}{j} \binom{\ell - d(x,x')}{k-j} = \mathcal{K}_k(d(x,x'); \ell, \alpha).$$

It follows that

$$\nu_k = \alpha^{-\ell} \sum_{x,x'} K_{x,x'} \mathcal{K}_k(d(x,x'); \ell, \alpha).$$

Next we compute the expected variance in $f$ given by

$$\mathbb{E}[f^T f] = \mathbb{E}\left[\sum_x f(x)^2\right] = \sum_x \mathbb{E}[f(x)^2] = \sum_x \mathrm{Var}[f(x)] = \sum_x K_{x,x} = \mathrm{tr}(K),$$

and thus

$$\bar{\nu}_k = \frac{\nu_k}{\mathbb{E}[f^T f]} = \frac{\sum_{x,x'} K_{x,x'} \mathcal{K}_k(d(x,x');\ell,\alpha)}{\operatorname{tr}(K)}. \tag{41}$$

□

**Lemma 2.** *Let $f \sim N(0, K)$, and let $K$ be a product kernel, $K_{x,x'} = \prod_p K^p_{x_p,x'_p}$ where $x_p$ denotes the character at position $p$ in sequence $x$. Then*

$$\nu_U = \alpha^{-\ell} \prod_{p\in U} (\alpha \operatorname{tr}(K^p) - \operatorname{sum}(K^p)) \prod_{p\notin U} \operatorname{sum}(K^p),$$

*where* tr *denotes the trace of the matrix and sum denotes the sum of all values in the matrix.*

*Proof.* Lemma 1 implies

$$\nu_U = \alpha^{-\ell} \sum_{x,x'} \left( \prod_{\substack{p\in U\\ x_p = x'_p}} (\alpha - 1) \prod_{\substack{p\in U\\ x_p \neq x'_p}} (-1) \prod_{p\in[\ell]} K^p_{x_p,x'_p} \right). \tag{42}$$

We consider how each position contributes to the summand for a pair of sequences and use this to rewrite this sum over all sequence pairs as a product over positions where each term in the expansion of the product equals the summand for one pair of sequences.

- If $p \in U$ and $x_p = x'_p = c$, the position $p$ contributes $(\alpha - 1)K^p_{c,c}$ to the summand.
- If $p \in U$, $x_p = c$ and $x'_p = c'$ where $c \neq c'$, the position $p$ contributes $-1K^p_{c,c'}$ to the summand.
- If $p \notin U$, $x_p = c$ and $x'_p = c'$, the position $p$ contributes $K^p_{c,c'}$ to the summand.

Note each term in the expansion of

$$\alpha^{-\ell} \prod_{p\in U} \left( (\alpha - 1) \sum_c K^p_{c,c} - \sum_{c\neq c'} K^p_{c,c'} \right) \prod_{p\notin U} \left( \sum_{c,c'} K^p_{c,c'} \right)$$

is equal to a summand for a pair of sequences $x, x'$ in Equation (42). It follows that

$$\nu_U = \alpha^{-\ell} \prod_{p\in U} \left( (\alpha - 1) \sum_c K^p_{c,c} - \sum_{c\neq c'} K^p_{c,c'} \right) \prod_{p\notin U} \sum_{c,c'} K^p_{c,c'} = \alpha^{-\ell} \prod_{p\in U} (\alpha \operatorname{tr}(K^p) - \operatorname{sum}(K^p)) \prod_{p\notin U} \operatorname{sum}(K^p).$$

□

*Proof.* (of Theorem 2.) The orthogonality of the subspaces $V_U$ implies that $\nu_k = \sum_{U:|U|=k} \nu_U$ and $\mathbb{E}[f^T f] = \sum_{k=0}^{\ell} \nu_k$. It follows from Lemma 2 that

$$\nu_k = \left( \alpha^{-\ell} \prod_{p\in[\ell]} \operatorname{sum}(K^p) \right) e_k \left( \frac{\alpha \operatorname{tr}(K^1)}{\operatorname{sum}(K^1)} - 1, \ldots, \frac{\alpha \operatorname{tr}(K^\ell)}{\operatorname{sum}(K^\ell)} - 1 \right),$$

where $e_k$ denotes the elementary symmetric polynomial of order $k$, i.e. the sum of all $k$-way products. We compute the average value of $K^p$ under the assumption that the diagonal is one,

$$\zeta_p = \frac{1}{\alpha^2} \left( \alpha + \sum_{c\neq c'} K^p_{c,c'} \right).$$

Noting that $\text{sum}(K^p) = \alpha^2\zeta_p$, we obtain

$$\frac{\alpha\,\text{tr}(K^p)}{\text{sum}(K^p)} - 1 = \frac{1}{\zeta_p} - 1.$$

It follows that

$$\bar{\nu}_k = \frac{\nu_k}{\mathbb{E}[f^T f]} = \frac{\nu_k}{\sum_j \nu_j} = \frac{e_k\left(\frac{1}{\zeta_1} - 1, \ldots, \frac{1}{\zeta_\ell} - 1\right)}{\prod_{p=1}^{\ell}\left(1 + \left(\frac{1}{\zeta_p} - 1\right)\right)} = \left(\prod_{p=1}^{\ell} \zeta_p\right) e_k\left(\frac{1}{\zeta_1} - 1, \ldots, \frac{1}{\zeta_\ell} - 1\right).$$

□

*Proof.* (of Theorem 3). For ease of notation, let $c = \alpha^{-\ell}\left(\prod_{p=1}^{\ell}(\alpha - (\alpha-1)\delta_p)\right)$. Note that $\bar{\nu}_k = ce_k\left(\frac{(\alpha-1)\delta_1}{\alpha-(\alpha-1)\delta_1}, \ldots, \frac{(\alpha-1)\delta_\ell}{\alpha-(\alpha-1)\delta_\ell}\right)$ if and only if

$$\sum_{k=0}^{\ell}(-1)^k \bar{\nu}_k x^{\ell-k} = c\prod_{p=1}^{\ell}\left(x - \frac{(\alpha-1)\delta_p}{\alpha - (\alpha-1)\delta_p}\right).$$

For the connectedness model to be well-defined (e.g. positive definite), each $\delta_p \in (0, 1 + \frac{1}{\alpha-1})$. It follows that for a set of $\delta_p$ that define a valid connectedness model,

$$\frac{(\alpha-1)\delta_p}{\alpha - (\alpha-1)\delta_p} \in (0, \infty)$$

for each $p$ meaning the polynomial has $\ell$ positive roots.

Next assume that the polynomial has $\ell$ positive roots. For each root $r_p$ we can solve the equation $\frac{(\alpha-1)\delta_p}{\alpha-(\alpha-1)\delta_p} = r_p$ and arrive at a $\delta_p \in (0, 1 + \frac{1}{\alpha-1})$. This set of $\delta_p$'s defines a connectedness model with the desired vector of normalized variance components.

□

#### Special Section References


[1] Carl Edward Rasmussen and Christopher K I Williams. *Gaussian processes for machine learning.* MIT Press, 2006.

[2] Juannan Zhou, Mandy S Wong, Wei-Chia Chen, Adrian R Krainer, Justin B Kinney, and David M McCandlish. "Higher-order epistasis and phenotypic prediction". In: *Proceedings of the National Academy of Sciences* 119.39 (2022), e2204233119.

[3] Philip A Romero, Andreas Krause, and Frances H Arnold. "Navigating the protein fitness landscape with Gaussian processes". In: *Proc. Natl. Acad. Sci. U.S.A.* 110.3 (2013), E193–E201.

[4] Luca Ferretti, Benjamin Schmiegelt, Daniel Weinreich, Atsushi Yamauchi, Yutaka Kobayashi, Fumio Tajima, and Guillaume Achaz. "Measuring epistasis in fitness landscapes: The correlation of fitness effects of mutations". In: *J. Theor. Biol.* 396 (2016), pp. 132–143.

[5] Claudia Bank, Sebastian Matuszewski, Ryan T Hietpas, and Jeffrey D Jensen. "On the (un)predictability of a large intragenic fitness landscape". In: *Proc. Natl. Acad. Sci. U.S.A.* 113.49 (2016), pp. 14085–14090. DOI: 10.1073/pnas.1612676113.

[6] Gautam Reddy and Michael M Desai. "Global epistasis emerges from a generic model of a complex trait". In: *Elife* 10 (2021), e64740.

[7] Carlos Martí-Gómez, Juannan Zhou, Wei-Chia Chen, Justin B. Kinney, and David M. McCandlish. "Inference and visualization of complex genotype-phenotype maps with *gpmap-tools*". en. In: *bioRxiv* (Mar. 2025).

#### Supplementary Figures

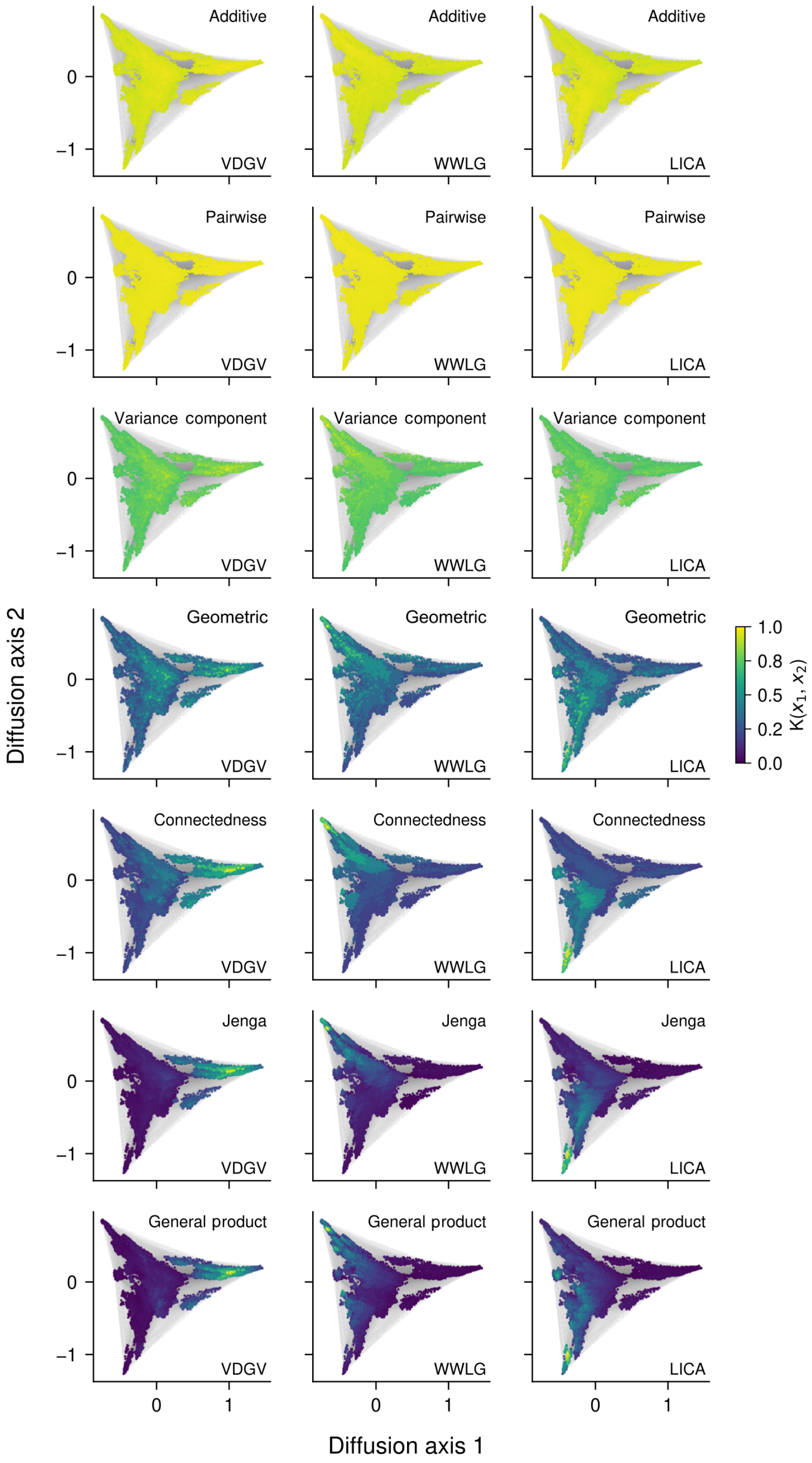


Figure S1: Low dimensional representation of the GB1 genotype-phenotype map inferred under the Jenga model using the complete dataset, where each dot represents a genotype, lines between them represent single point mutations, and squared distances optimally approximate the commute times between genotypes under a weak mutation evolutionary model [74, 93]. Genotypes are colored according to their correlations under the indicated prior with the specific sequences at the bottom right corner of each panel. These visualizations highlight how the more expressive priors are able to capture more detailed information about the three main fitness peaks of this functional landscape [58].

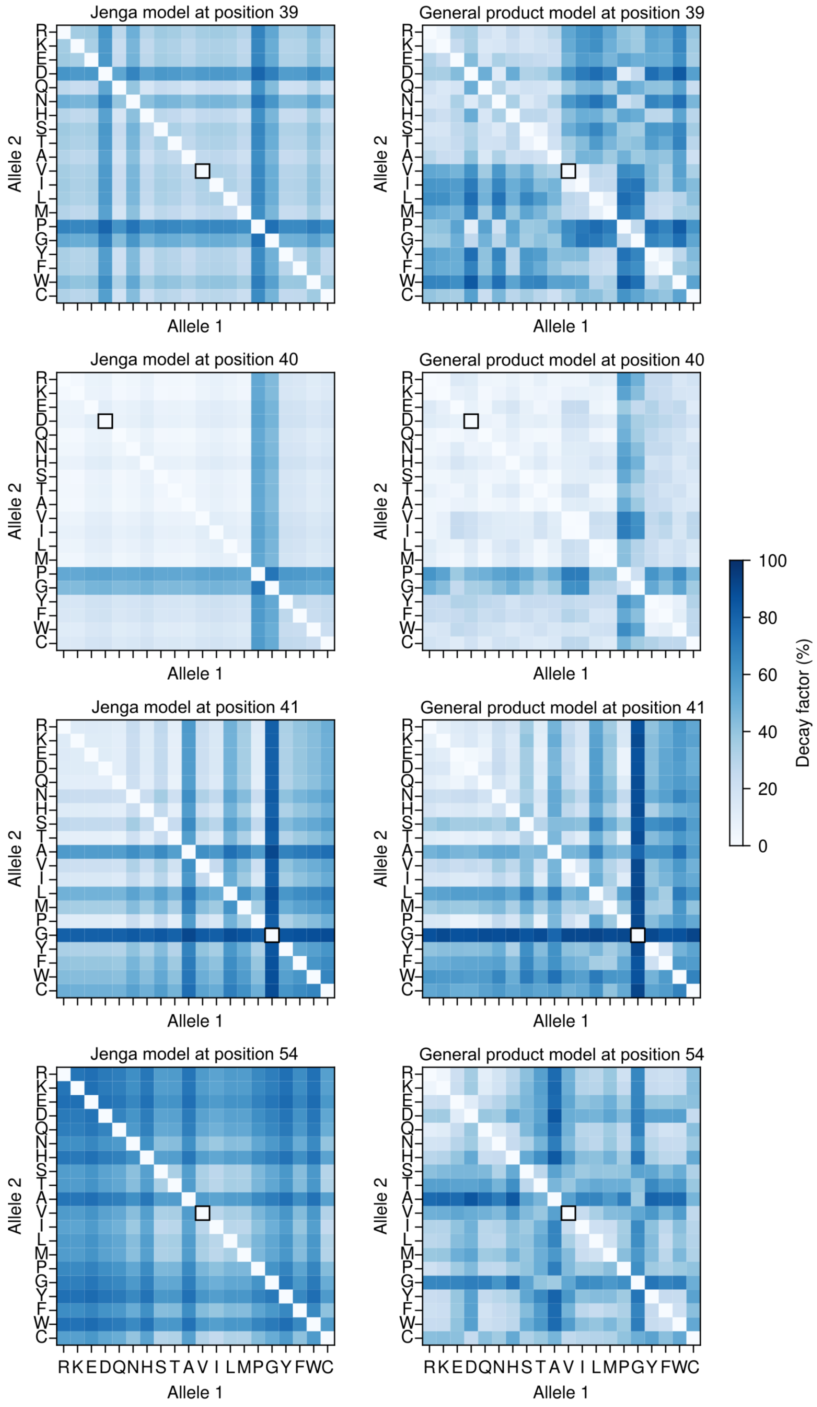


Figure S2: Comparison of the mutation-specific decay factors inferred under the Jenga (left) and general product (right) models across the four sites of the GB1 binding protein landscape.

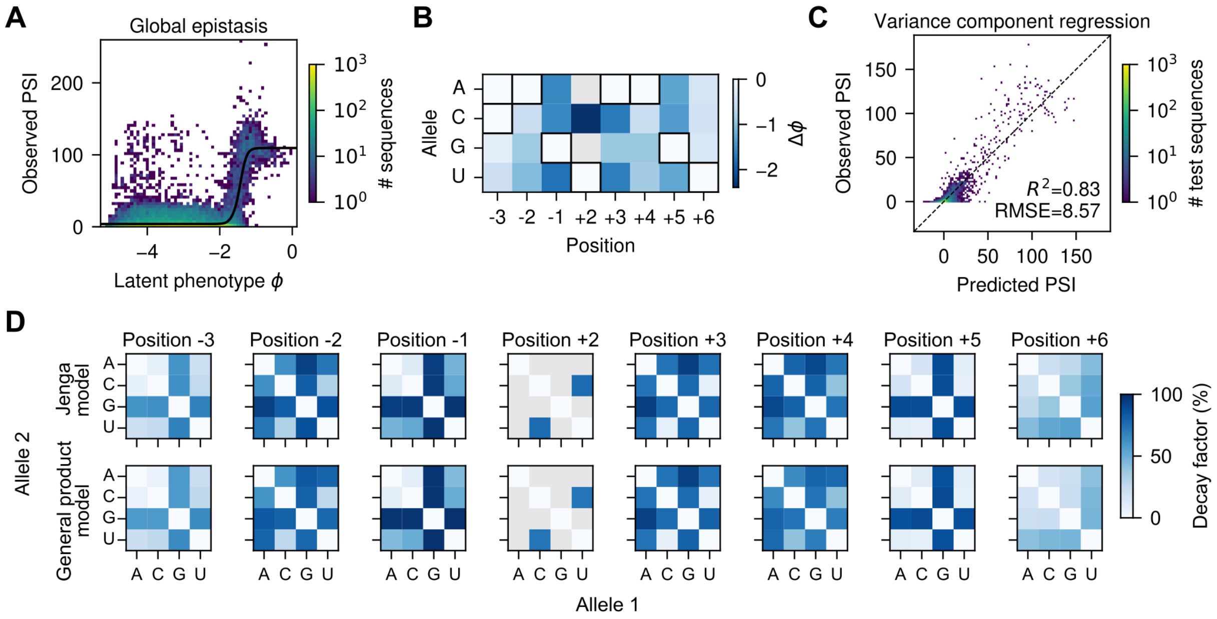


Figure S3: Models of sequence-function relationships for the 5′ splice site in the SMN1 exon 7 context. (A) Comparison of the observed PSI with the inferred latent phenotype under an additive global epistasis model using MAVE-NN [61]. The black line represents the learned non-linear relationships between the latent phenotype and the measured PSI. (B) Estimated single point mutational effects under the global epistasis model. Latent phenotype and mutational effects are normalized so that the maximum value of the latent phenotype is 0 and the distribution of mutational effects has a standard deviation of 1 (A,B). (C) Comparison of the observed and predicted PSI under the variance component regression model showing a non-linear dependency of the residuals on the predicted values. PSI: Percent Spliced In (A-C). (D) Comparison of the mutation-specific decay factors inferred under the Jenga (top) and general product (bottom) models across all sites of the SMN1 5′ splice site sequence landscape.

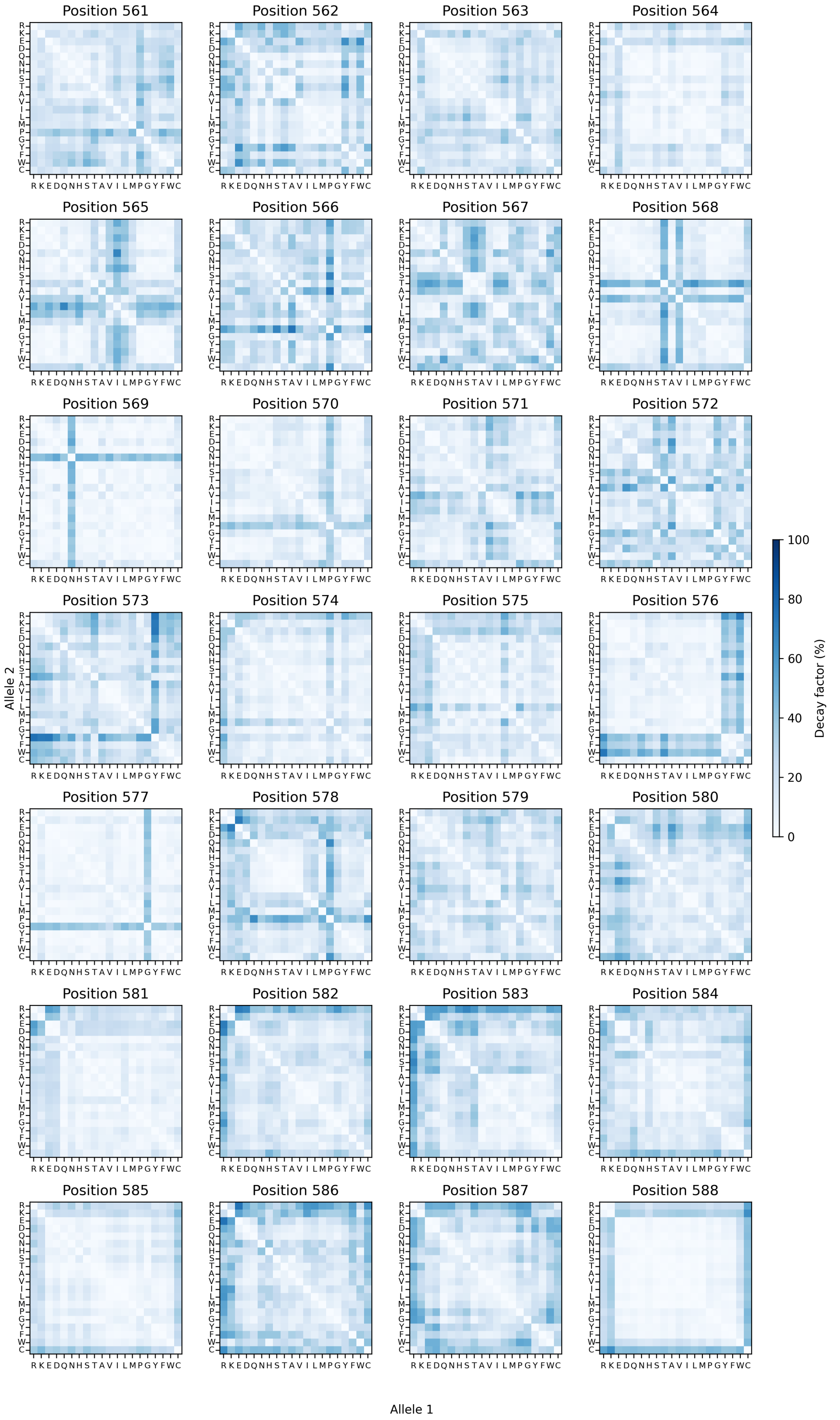


Figure S4: Mutation-specific decay factors inferred under the general product model for every site of the AAV2 capsid protein dataset.

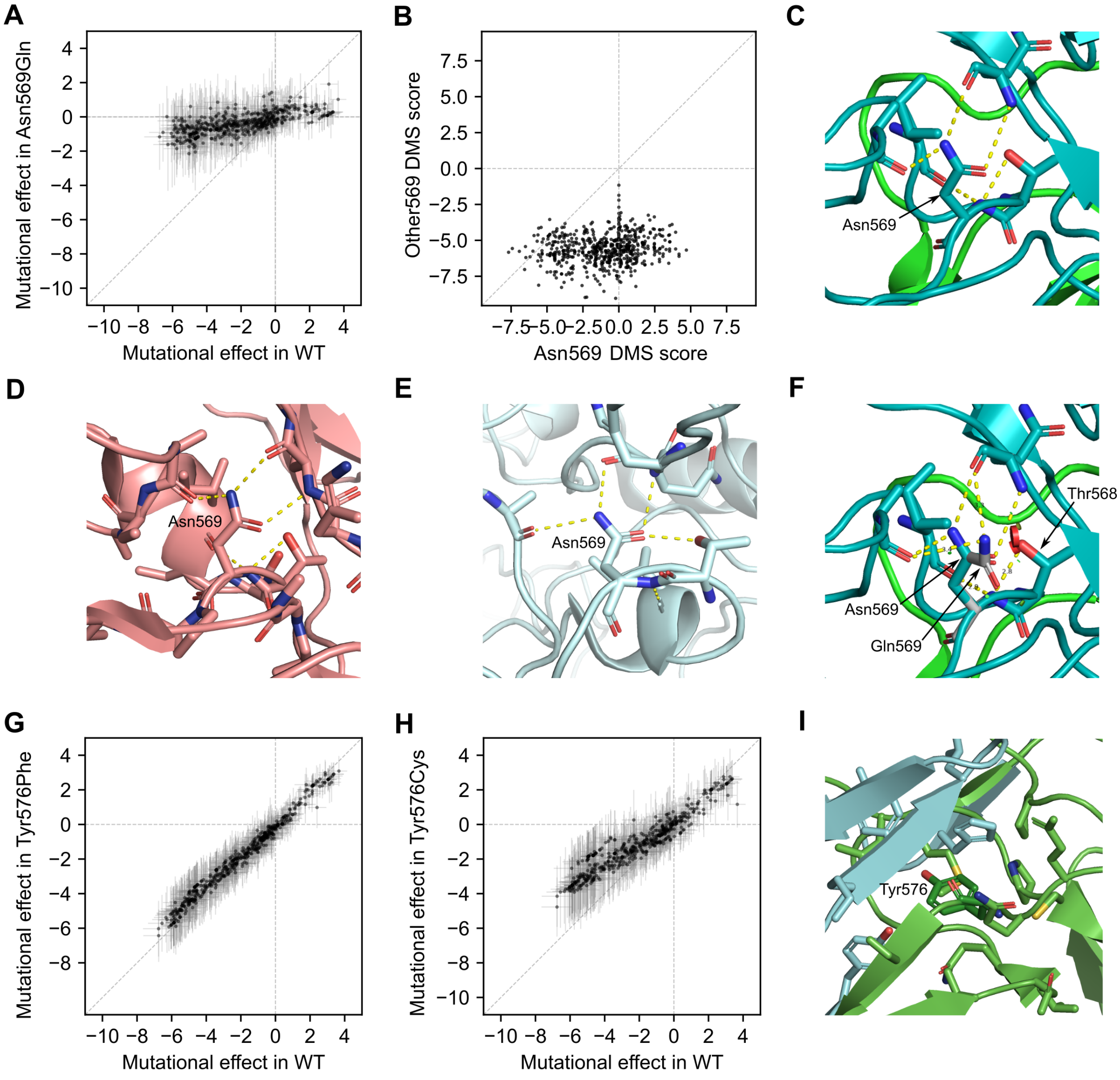


Figure S5: Pervasive effects of mutations at position 569 and 576 on the effects of other mutations in the AAV2 capsid protein. (A) Comparison of the estimated mutational effects under the Jenga model in the wild-type (WT) context and in the presence of the most conservative substitution at position 569 from Asn to Gln. This comparison shows the systematic reduction in the effects of all other mutations. (B) Scatterplot of the experimentally determined DMS scores of sequences containing Asn at position 569 compared the same sequence with any other amino acid at that position. This shows that genetic variants with effects in the presence of Asn no longer have an effect in the presence of other amino acids at 569. (C-E) AAV2 capsid protein structure showing that Asn lateral chain at position 569 forms 3 hydrogen bonds with the backbone at positions 522 and 607 within the same monomer in AAV2 (PDB:8FZ0) (C) and AAV3 (PDB:8A9U) (D), and 4 hydrogen bonds with backbone of 522 and 607, and the Thr at 568 in AAV1 (PDB:8FQ4) (E). (F) AAV2 capsid protein structure showing the orientation of Gln at position 569 compared to Asn as predicted by Pymol. Despite being able to preserve the stabilizing hydrogen bonds, the larger size of Gln would imply a steric clash with Thr568 illustrated by the red disc. (G,H) Estimated mutational effects under the Jenga model in the wild-type (WT) context and in the presence of another aromatic residue (Phe, G) and non-aromatic residue (Cys, H) at position 576 showing their impact on the effects of all other mutations. (I) Structure showing that Tyr at position 576 is located in a hydrophobic pocket involving two of the monomers in the assembled AAV2 capsid (PDB:8FZ0). Different capsid monomers are represented in different colors (C, D, E, F, I)

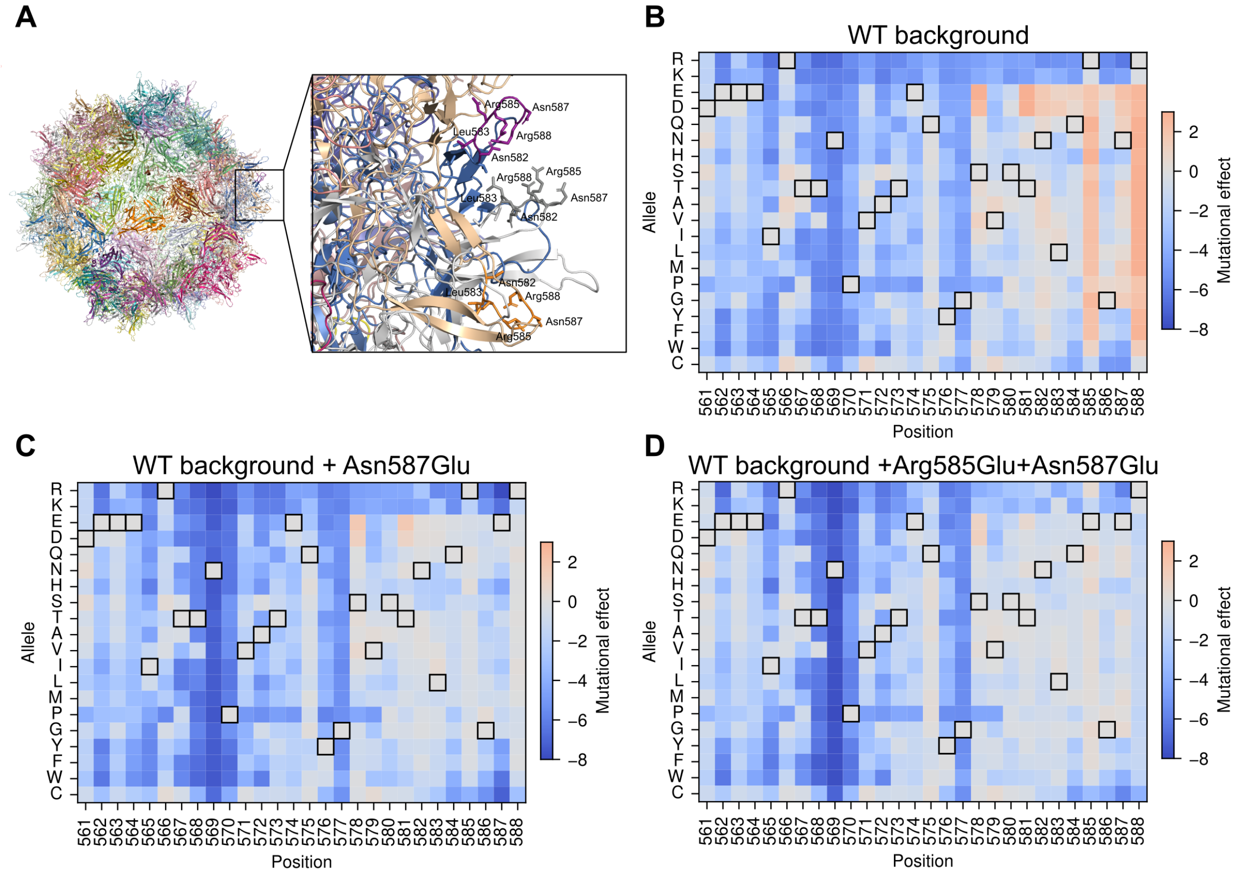


Figure S6: AAV2 capsid packaging requires an intermediate charge at the exposed loop comprising residues 579-588. (A) Structure of the empty AAV capsid highlighting the 579-588 protein region across all monomers and zoom into the structure of those loops at 3 adjacent monomers, represented with different colors. (B-D) Estimated mutational effects under the Jenga model for the wild-type background (B) and when introducing negative charges in the 579-588 region i.e. Asn587Glu alone (C) or together with Arg585Glu (D), illustrating the change in the sign of mutational effects that decrease positive charge in this region

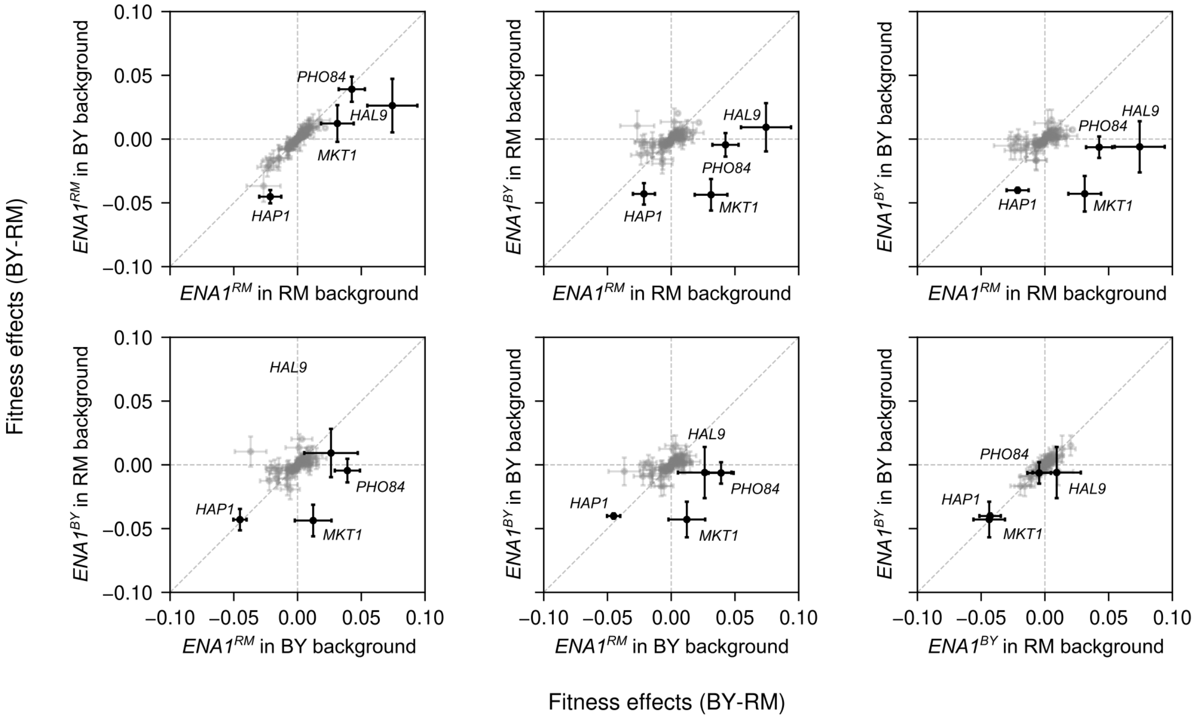


Figure S7: *ENA1* reshapes the genotype–phenotype map of yeast fitness under high $Li^{+}$ conditions. Estimated fitness effects (from RM to BY alleles) obtained using connectedness regression are compared across genetic backgrounds differing at the ENA1 locus, as well as across the overall genetic background (all other loci with BY or RM alleles). The results highlight the strong context dependence of specific mutations. Error bars represent the standard deviation of the posterior distribution for each fitness effect.
